# Marked regional glial heterogeneity in the human white matter of the central nervous system

**DOI:** 10.1101/2022.03.22.485367

**Authors:** Luise A. Seeker, Nadine Bestard-Cuche, Sarah Jäkel, Nina-Lydia Kazakou, Sunniva M. K. Bøstrand, Alastair M. Kilpatrick, David Van Bruggen, Mukund Kabbe, Fabio Baldivia Pohl, Zahra Moslehi, Neil C. Henderson, Catalina A. Vallejos, Gioele La Manno, Gonçalo Castelo-Branco, Anna Williams

**Affiliations:** Centre for Regenerative Medicine, Institute for Regeneration and Repair, Edinburgh Bioquarter, University of Edinburgh, 5 Little France Drive, Edinburgh EH16 4UU; Institute for Stroke and Dementia Research, Klinikum der Universität München, Ludwig-Maximilians-Universität, Munich, Germany; Munich Cluster for Systems Neurology (SyNergy), Munich, Germany; Laboratory of Molecular Neurobiology, Department of Medical Biochemistry and Biophysics, Karolinska Institutet, 171 77 Stockholm, Sweden; Laboratory of Neurodevelopmental Systems Biology, Brain Mind Institute, School of Life Sciences, École Polytechnique Fédérale de Lausanne (EPFL), 1015 Lausanne, Switzerland; Centre for Inflammation Research, The Queen’s Medical Research Institute, Edinburgh BioQuarter, University of Edinburgh, Edinburgh, UK; MRC Human Genetics Unit, Institute of Genetics and Cancer, University of Edinburgh, Western General Hospital, Edinburgh, EH4 2XU, UK; The Alan Turing Institute, 96 Euston Road, London, NW1 2DB, UK; Ming Wai Lau Centre for Reparative Medicine, Stockholm node, Karolinska Institutet, 171 77 Stockholm, Sweden

## Abstract

The myelinated white matter tracts of the central nervous system (CNS) are essential for fast transmission of electrical impulses and are commonly affected in neurodegenerative diseases. However, these often uniquely human diseases differentially affect white matter regions, at various ages and between males and females, and we hypothesised that this is secondary to physiological variation in white matter glia with region, age and sex. Using single nucleus RNA sequencing of healthy human post-mortem samples, we find marked glial heterogeneity with tissue region (primary motor cortex, cerebellum, cervical spinal cord), with tissue-specific cell populations of oligodendrocyte precursor cells and astrocytes, and a spinal cord-enriched oligodendrocyte type that appears human-specific. Spinal cord microglia but not astrocytes show a more activated phenotype compared to brain. These regional effects, with additional differentially expressed genes with age and sex in all glial lineages, help explain pathological patterns of disease – essential knowledge for therapeutic strategies.

## Introduction

The cells of the white matter (WM) of the human central nervous system (CNS) are mostly glia. These cells (oligoden-droglia - oligodendrocytes and their oligodendrocyte precursor cells (OPCs), astrocytes and microglia) are of crucial importance for the physiology of the CNS but are also key players in the pathologies of demyelinating, neurodegenerative and neuropsychiatric disorders (reviewed in^1^). Some of these diseases have pathological biases to CNS region, are more common in one sex and increase in incidence with age. Animal model data suggest that functional differences in glia may causally underpin this observed variation in pathology. In mouse, there is morphological and functional diversity in oligodendroglia in different white matter regions^2, 3^ linked to differences in tissue environment^4^, axon calibre^5^, extra cellular matrix (ECM) stiffness^6^, and intrinsic factors related to their developmental origin^57^. Aged rodent OPCs proliferate and differentiate slower^8–10^, multiple sclerosis (MS)-like WM demyelinated lesions in aged mice remyelinate less efficiently^11^, ageing rhesus monkeys show more pro-inflammatory activated, phagocytic microglia^12^ and in humans, age is the main risk factor for more severe progressive MS^13^. Sex dimorphism in neurodegenerative disease susceptibility suggests that either gonosomal genes or gonadal hormones may also affect the function of CNS^14–16^, detectable at the individual cell level: cultured female neonatal rat OPCs show more proliferation, migration and less differentiation than males^17^, and there are more anti-inflammatory maintenance microglia in adult female mice^18^.

However, despite these data in animal models and their presumed link to disease, WM glial regional, age and sex diversity is not well understood in humans. Single-nucleus transcriptomics data from post-mortem human samples have been very successfully used to study cellular changes in development^19–24^, or disease^25–28^ and we used this technology to address our hypothesis that glial diversity is an important mediator of region, age and sex differences in the normal human CNS.

Here, we performed a transcriptional characterisation of human WM cells of healthy post-mortem samples from three different CNS regions: Primary motor cortex (Brodmann area 4; BA4), cerebellum (CB) and cervical spinal cord (CSC), from donors of two different age groups (30 – 45y and 60 – 75y) and both sexes. We found significant regional heterogeneity with region-specific populations of oligodendrocyte precursor cells (OPCs) and astrocytes, and spinal cord-enriched populations of oligodendrocytes and microglia. Transcriptional variation with age and sex were also identified, with most marked effects in OPCs, microglia and astrocytes. We discuss potential causes and the expected impact of these differences on health and targeting therapies in disease, and provide an open-source atlas for researchers as a comparator for pathologies.

## Results

### Description and annotation of the complete dataset

We used WM from three anatomical CNS sites selected for marked structural differences: BA4, CB and CSC. We analysed these samples over a cohort of 20 British Caucasian donors (60 samples in total) (Figure 1a, Extended Data Table 1). Donors equally represented both sexes and two different age groups “young adults” (30-45 y, 5 males and 5 females) and “old adults” (60-75 y, 5 males and 5 females). After strict sample, cluster and nucleus quality control (see methods, Extended Data Figure 1, Table S1) we retained 48 samples and 48,104 nuclei (mean number of genes per nucleus: 1853, mean number of UMIs per nucleus: 5450). A first level cluster analysis and marker inspection revealed all expected major cell types: excitatory neurons (as marked by *SNAP25, SLC17A7*), inhibitory neurons (*SNAP25, GAD1*), reelin-positive neurons (*SNAP25, RELN*), astrocytes (*GJA1, GFAP*), microglia and macrophages (*CD74, P2RY12*), endothelial cells and pericytes (*CLDN5, NOTCH3*), oligodendrocytes (*PLP1, CNP*), their precursor cells (*PDGFRA, PTPRZ1*) and immune cells (*HLA-A, PTPRC*) (Figure 1b–c). We subsetted the data for all main cell lineages and re-clustered each resulting dataset allowing finer distinction of cellular states, identifying 11 oligodendroglia, 11 astrocyte, 6 microglia and macrophage, 11 vascular and 18 neuronal clusters that expressed marker genes (Figure 1b, Table S2).

**Figure 1.**
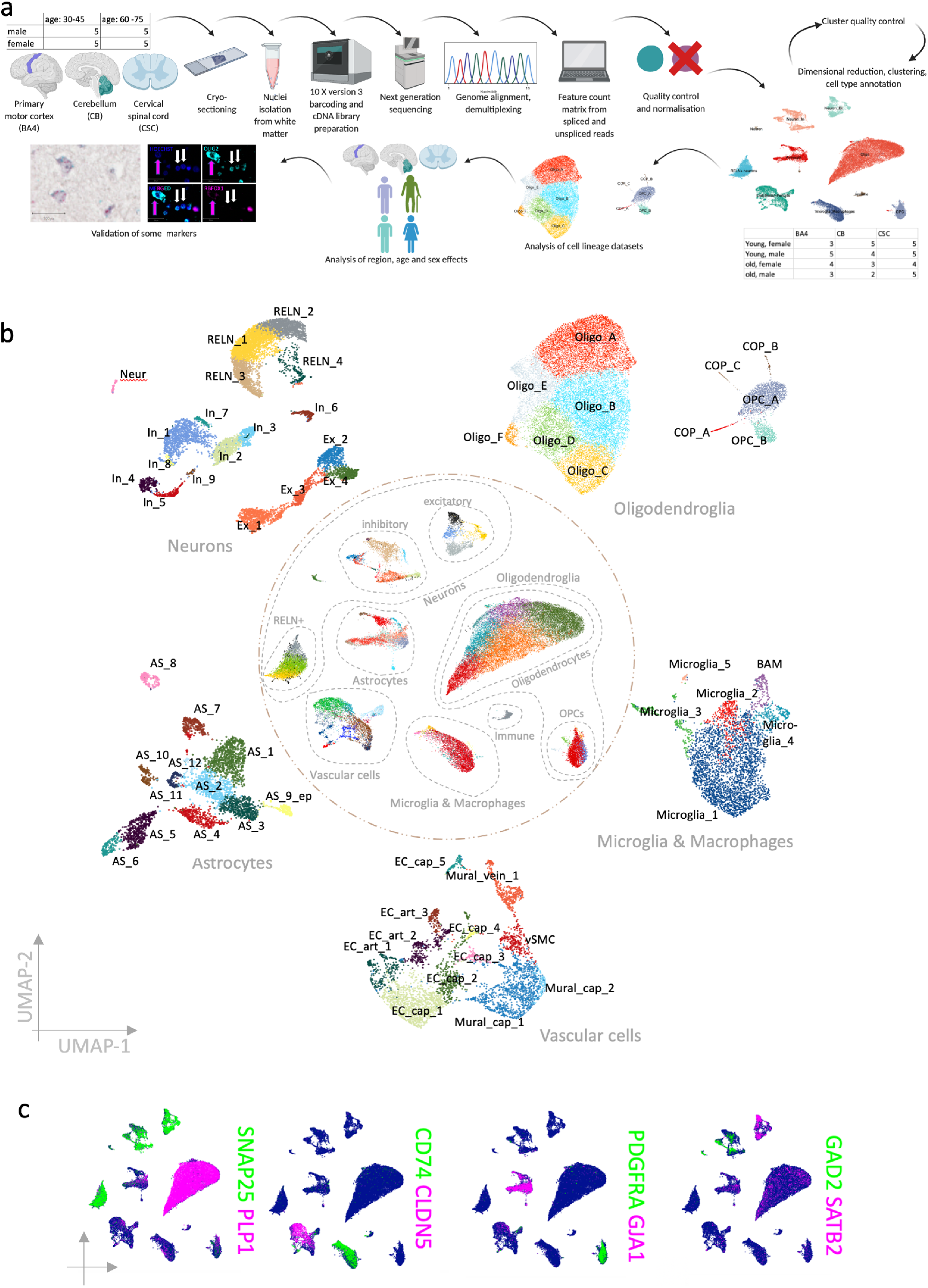
Complete dataset. a) Schematic of the workflow. b) UMAP representation of the complete dataset in the centre and cell lineage datasets at the circumference. c) Selection of canonical cell lineage marker gene expression in the complete dataset.

### Oligodendroglia are transcriptionally heterogeneous

We first analysed oligodendroglia, the most abundant cell lineage in the human WM. The cluster analysis revealed six oligodendrocyte states (positive for myelin genes such as *PLP1*, *MBP*, *MAG*), two OPC states (positive for *PDGFRA*, *CSPG4* and *BCAN*) and three committed oligodendrocyte precursor (COP) cell states (positive for *GPR17*, *GPC5* or *GAP43*), confirming that oligodendroglia are heterogeneous in the healthy adult human CNS (Figure 2a–c).

**Figure 2.**
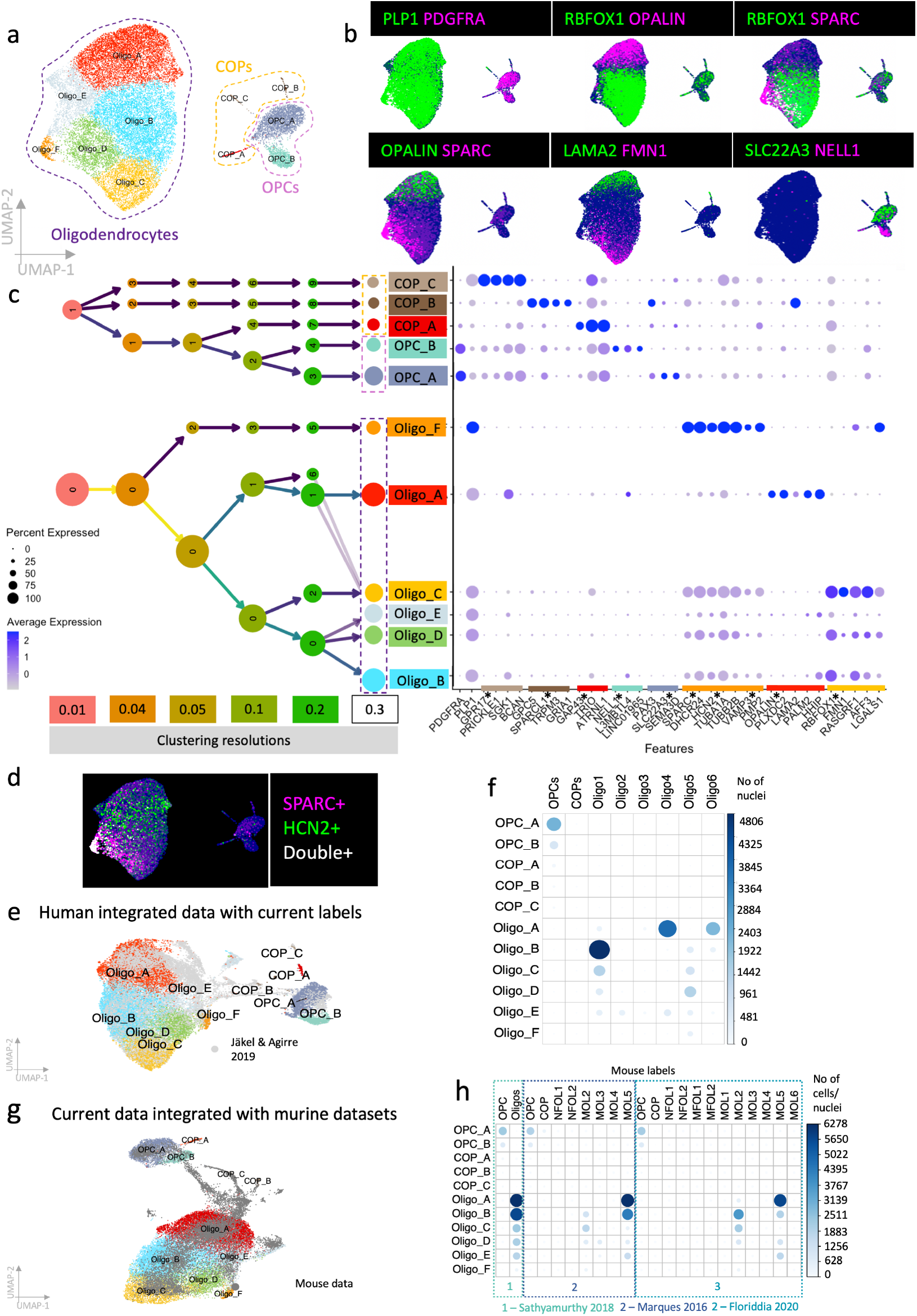
Oligodendroglia dataset. a) finalised clustering, b) expression of lineage marker and cluster marker genes, c) Redistribution of nuclei to clusters depending on different clustering resolutions (0.01 – 0.3) shows relationship of clusters. On the left, cluster 0 represents oligodendrocytes, cluster 1 represents OPCs. Dot plot shows marker genes for oligodendroglia clusters, with * indicating those mentioned in text. d) Co-expression of *SPARC* and *HCN2*. e) CCA integration of current dataset with previously published human dataset25 (current cluster labels include letters to clearly distinguish them from previous cluster labels). f) Correlation of current and predicted human labels25 (label transfer). g) CCA integration with three murine oligodendroglia datasets2, 3, 40. h) Correlation between human labels and predicted mouse labels (label transfer).

*RBFOX1* is a marker for the oligodendrocyte clusters Oligo_B-E, which subcluster on the basis of expression of other genes: Oligo_B: *AFF3*+ *LGALS1*- *FMN1*-; Oligo_C: *FMN1*; Oligo_D: *LGALS1*+ *FMN1*- *SPARC*-; Oligo_E: *HHIP*+ *OPALIN*- *AFF3*- (2b,c), although Oligo_E has no exclusively enriched marker genes, and so may be the default oligodendrocyte population from which other oligodendrocyte clusters differ by additive expression of specific genes (2c, Table S2). Oligo_A and F are *RBFOX1*-negative and strikingly different, Oligo_A expressing *OPALIN*, *PLXDC2*, *LAMA2* and *PALM2*, and Oligo_F expressing *SPARC*, *DHCR24*, *TUBA1A*, *TUBB2B*, *PMP2* (2c). Gene ontology (GO) analysis for Oligo_A revealed 52 enriched pathways (Table S3) including those important for interaction with neurons, such as axon development, regulation of neuronal projections and GTPases mediated signal transmission (underpinned by genes such as *STARD13*, *AUTS2* and *ANK3*) and the expression of *ANK3* and *OPALIN* suggests a link with paranodal development and/or maintenance^29, 30^ (Extended Data Figure 2a, Table S3). Oligo_F expresses the novel oligodendrocyte marker *SPARC* and is mostly found in CSC (see below). GO analysis for Oligo_F markers revealed 405 statistically enriched terms, including 19 terms based on genes such as *MIF* and *GSTP1* suggesting an immune-related function. However, these are different from the immune oligodendroglia we previously described^31^, as these oligodendrocytes do not express MHC-2 genes, *CD74* nor *CTSS*. Oligo_F shows enriched genes related to sterol, cholesterol and myelin production (Extended Data Figure 2b–c, Table S3), and increased expression of the Snare protein gene *VAMP3* implicated in myelin sheath production^32^ suggesting increased myelin production and/or maintenance. Oligo_F also express the highest levels of *HCN2* which is important for the formation of longer myelin sheaths in mice^33^ (Figure 2d).

We found two OPC clusters, OPC_A, expressing *PAX3* and *SLC22A3*, suggesting a dorsal origin^34^ and OPC_B, expressing *NELL1* (Figure 2a–c, Table S2). COP_C expresses traditional COP markers such as *GPR17*, but the other COP clusters (albeit very limited in number in this adult dataset) are low in *GPR17* expression (2c). COP_B expresses genes of the astrocyte lineage including *SPARCL1* (2c), which has previously been detected in human foetal astrocyte-oligodendrocyte precursor cells^22^ and pre-OPCs^34^. As *SPARCL1* and *SPARC* (a marker for Oligo_F) are highly related, and there are more COP_B and Oligo_F cells in the CSC compared to the other tissues (see below), this raises the question as to whether COP_Bs give rise to Oligo_F cells.

### No linear differentiation trajectory is identifiable in healthy adult oligodendroglia

To test this hypothesis and look for evidence for a differentiation trajectory in the oligodendroglia, we used different trajectory inference (TI) methods both inside of Dynverse^35^(Angle, Scorpius, PAGA Tree) and standalone (Monocle^36^, Slingshot^37^, scVelo^38^) arguing that a convincing trajectory should be confirmed by several methods. However, we obtained conflicting results with disagreement as to the immature oligodendrocyte cluster: Oligo_B (Slingshot^37^, PAGA Tree) or Oligo_A (Monocle^36^, Angle, Scorpius) or an inverted trajectory using scVelo^38^, if performed on all oligodendroglia, or separated by region, apparently driven by methodological underlying assumptions and restrictions (linear, number of possible forks, circular trajectories) and not by oligodendroglial biology (Extended Data Figure 3). There may be little on-going oligodendroglial differentiation in our non-diseased adult human CNS dataset, supported by no evidence for cycling OPCs (negative for cell cycling markers e.g. *MKI67* and with Seurat’s cell cycle annotation algorithm^39^) and sparse COPs (intermediate cells), suggesting possible fixed cell endstates or dynamic switching between multiple states in response to environmental changes which TI methods cannot identify.

### Integrated analysis of oligodendroglia datasets reveals a human-specific oligodendrocyte subtype

To determine how our 11 oligodendroglial subclusters relate to published datasets, we transferred the labels of our study by Jäkel & Agirre et al. (2019)^25^ which included human samples of MS subcortical white matter and disease-free controls onto our dataset and then integrated both datasets (Figure 2e, Extended Data Figure 4). The predicted labels fitted well with our cluster labels, particularly for cluster Oligo_B which mostly corresponds to cluster Oligo1 and Oligo_A which corresponds best to Oligo6 and Oligo4. Clusters Oligo2 and Oligo3 that are enriched in MS^25^ are very rare in our current dataset, which confirms that those cellular states are MS-selective (2e,f). OPCs in Jäkel & Agirre et al.^25^ are of mixed brain regions and correlate to both of our OPC states (2f). The predicted labels for all of our COPs are defined as OPCs in the Jäkel & Agirre dataset^25^, again suggesting the absence of later intermediate cells that stop expressing OPC genes in our adult dataset where there is no pathology. Oligo_F does not map well to any cluster which underlines its spinal cord specificity (2f).

To check whether there is an equivalent of Oligo_F in mouse spinal cord, we also integrated our oligodendroglia dataset with three murine datasets by Marques et al.^3^ (different CNS regions of juvenile mice), Sathyarmurthy et al.^40^ (adult mouse lumbar spinal cord) and Floriddia et al.^2^ (adult corpus callosum and spinal cord) (2g,h). Human and mouse data integrated well, with OPC_A, Oligo_A and Oligo_B correlating particularly well to mouse oligodendroglia. However, newly formed oligodendrocytes are absent from the human dataset and the *SPARC*-positive, *RBFOX1*-negative human cluster Oligo_F is missing in the mouse datasets, even in the adult mouse spinal cord samples, suggesting it may be human-specific (2g,h). There are proteomic differences between human and murine CNS myelin including the human-specific expression of Peripheral Myelin Protein 2 (*PMP2*)^41^, which is highly expressed in Oligo_F and therefore may contribute to this observed proteomic species differences. More intermediate cells between the state of OPCs and oligodendrocytes were found in the mouse data, particularly in the juvenile data (2g,h), consistent with our data suggesting that human adult normal CNS cells are not actively differentiating.

### Transcriptional differences among astrocytes reveal division of labour but no ‘activation’ in normal adult brain

Astrocytes contribute to the formation of the blood brain barrier (BBB), metabolically support axons, control extracellular neurotransmitters, ions and fluids, regulate blood flow and are important for the development and pruning of synapses (reviewed in^42^). Human grey matter (GM) astrocytes are described as transcriptionally heterogeneous^43–45^, but similar literature on human WM astrocytes is much sparser^25, 26^. Our analyses identified twelve astrocyte WM clusters (AS_1 – AS_12) with AS_9_ep representing transcriptionally closely related ependymal cells, all with distinct marker gene expression (Figure 1b, 3a–b, Table S2). GO analysis suggests that astrocyte functional roles are divided out, with two types (AS_1 and 2) expressing genes concerned with neuronal interactions, axon guidance and synapse organisation, three (AS_4, 6 and 12) related to synapses/synaptic vesicles, two populations (AS_8 and 11) express myelin genes, as previously described in human astrocytes^46^, AS_5 relate to ECM organisation, AS_7 to the BBB and AS_10 to cellular motility, signalling and histone modification. Ependymal cells (AS_9_ep) express motile cilia genes (*CFAP43, SPAG17, DNAH6*) in keeping with their function in promoting cerebrospinal fluid flow^47^ (Extended Data Figure 5a, Table S3).

**Figure 3.**
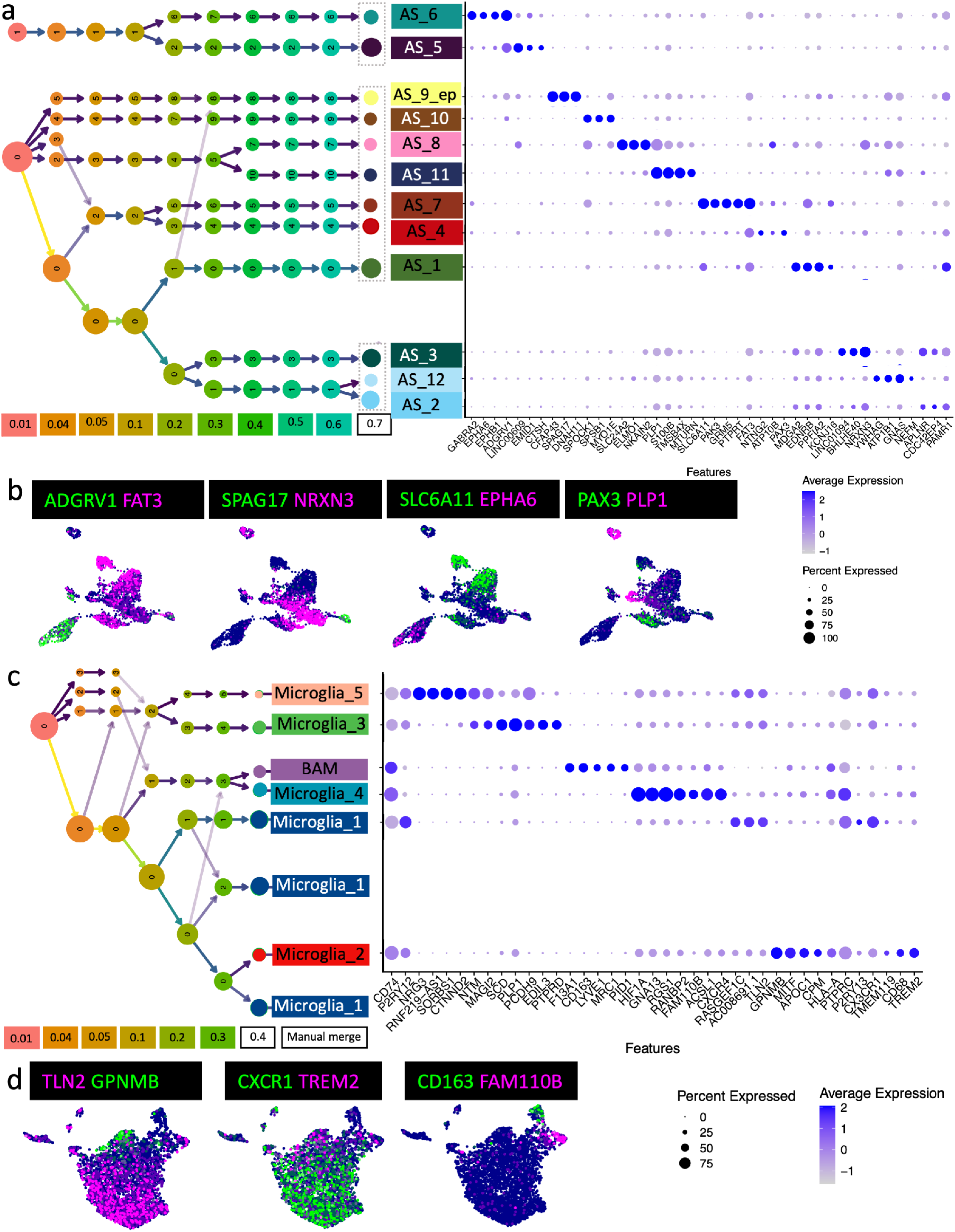
Astrocytes and microglia. a) astrocyte cluster tree visualises redistribution of nuclei to clusters based on increased cluster resolution. Dot plot shows the expression of selected cluster marker genes. b) Expression of selected cluster marker genes in the astrocyte lineage. c) Microglia cluster tree of cluster redistribution and dot plot of microglial cluster marker genes. d) Expression of selected marker genes for microglial clusters and activation.

Astrocytes change morphology and function upon injury or disease, initially categorised into a neurotoxic A1 type and a neuroprotective A2 type in mice^48, 49^, but now considered more of a spectrum^50, 51^, requiring combined markers to define the astrocyte activation state. Ageing may be interpreted as a chronic low-level injury reflected in astrocytes in old mice showing a more activated phenotype^52^, but we found no evidence of genes associated with astrocyte activation in our adult normal dataset, with no increase with age (Extended Data Figure 6).

### Activated microglia are present in normal adult CNS

We identified five microglia clusters (Microglia_1 – Microglia_5) and one cluster of border-associated macrophages (BAM – positive for *CD163, LYVE1, MRC1*) which express distinct marker genes (Figure 1b, 3c–d, Table S2), but no cells with the phenotype of infiltrating monocyte-derived macrophages, expected to be absent in healthy tissue. Similarly to astrocytes, microglia can shift from a homeostatic to an activated phenotype, characterised by proliferation, chemoattractant-mediated migration, changes in cellular morphology and phagocytosis of damaged cells^53, 54^. Microglia_1 (*RASGEF1C, AC008691.1, TLN2*) express markers and GO terms for a homeostatic phenotype (*P2RY13 and CX3CR1*) (Figure 3c, Extended Data Figure 7, Table S2, Table S3). Conversely, Microglia_2 (*GPNMB*) and Microglia_4 (*ACSL1, CXCR4*) are activated, based on their expression of the phagocytotic markers *CD68* and *TREM2* and their associated significant leukocyte activation and degranulation GO terms, in spite of being derived from healthy tissue (Extended Data Figure 7a, Table S2, Table S3). However, even in normal adult tissue, activated microglia are needed for regulation of neuronal survival, phagocytosis of apoptotic excess neurons life^55^, and pruning of synapses which is important for circuit refinement^56^. Microglia_3 and 5 express genes normally expressed by neurons or glia such as *NTM, MAGI2, SCD, PLP1, NRG3*, suggesting that these were present due to microglial engulfment of oligodendrocytes and/or synapses (Table S2, Table S3).

### Regional heterogeneity is marked in WM glia

We next focussed on how clusters varied depending on region. Compositional differences of all cell lineages with region were obvious from inspecting dimensionally reduced representations of the datasets (Figure 4a–c), but also confirmed (with measures of significance) using Milo^57^(Figure 4d). In oligodendroglia, we observed the most striking compositional difference with region in the OPC population: OPC_A was predominantly found in CB and CSC, whereas Oligo_B was more abundant in BA4 (Figure 4d). This reflects the different developmental origins of OPCs in those CNS regions^58^, as *PAX3*+ dorsal origin CSC OPCs also more highly express the *HOX* genes, important for craniocaudal and dorso-ventral development while BA4 OPCs express the anterior marker *FOXG1* (Figure 5a–e). We did not identify previously described cerebellum-specific OPCs that express *ORAOV1+LRP6+RCN2+*^59^ in the present study, perhaps as we only captured WM.

**Figure 4.**
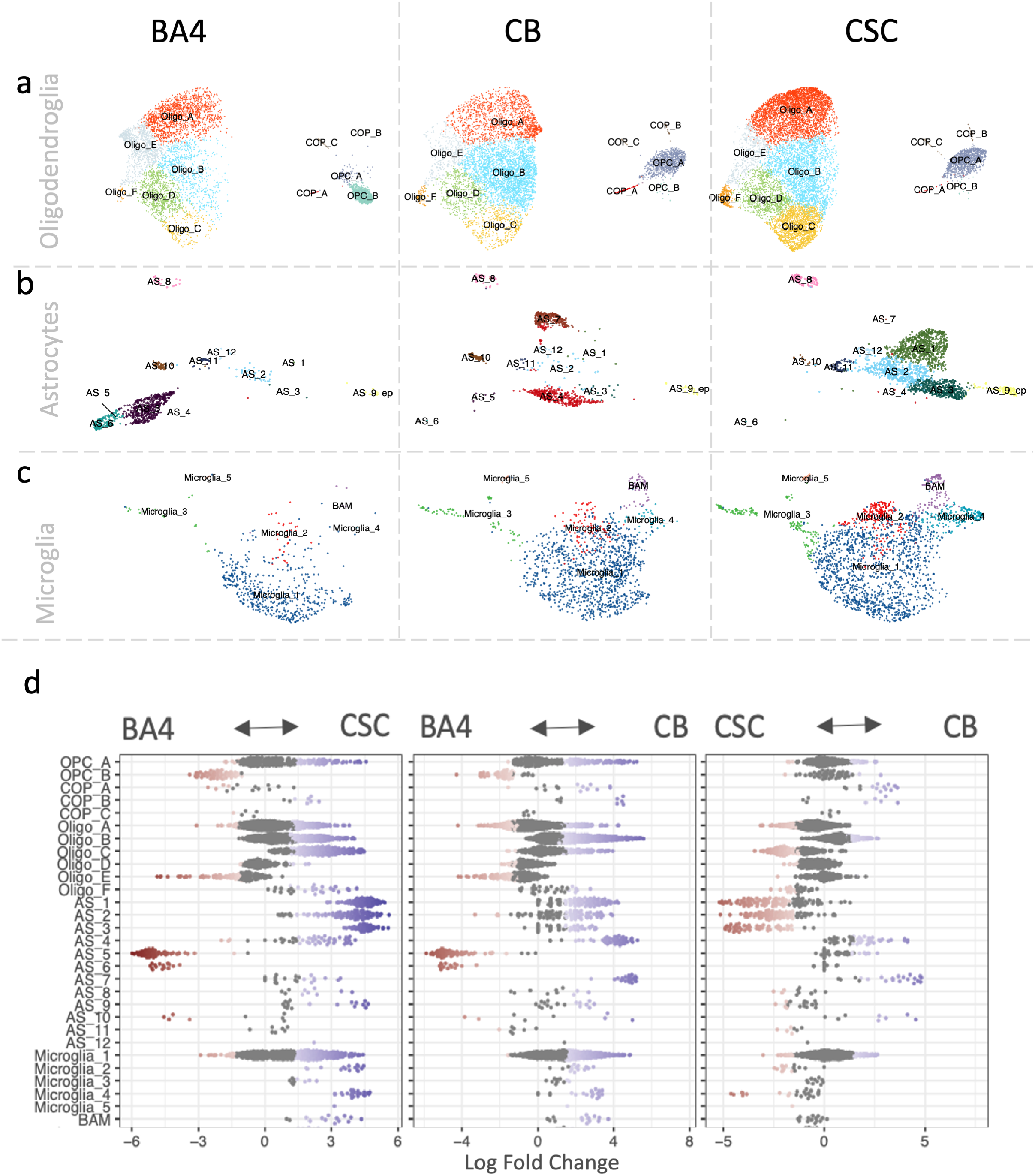
Glial compositional differences with tissue region. UMAP representation of a) oligodendroglia, b) astrocytes, c) microglia. d) Compositional analysis using Milo57 shows statistically significant compositional differences (blue and red). BA4 = white matter underlying motor cortex, CB = white matter of cerebellum, CSC = white matter of cervical spinal cord.

**Figure 5.**
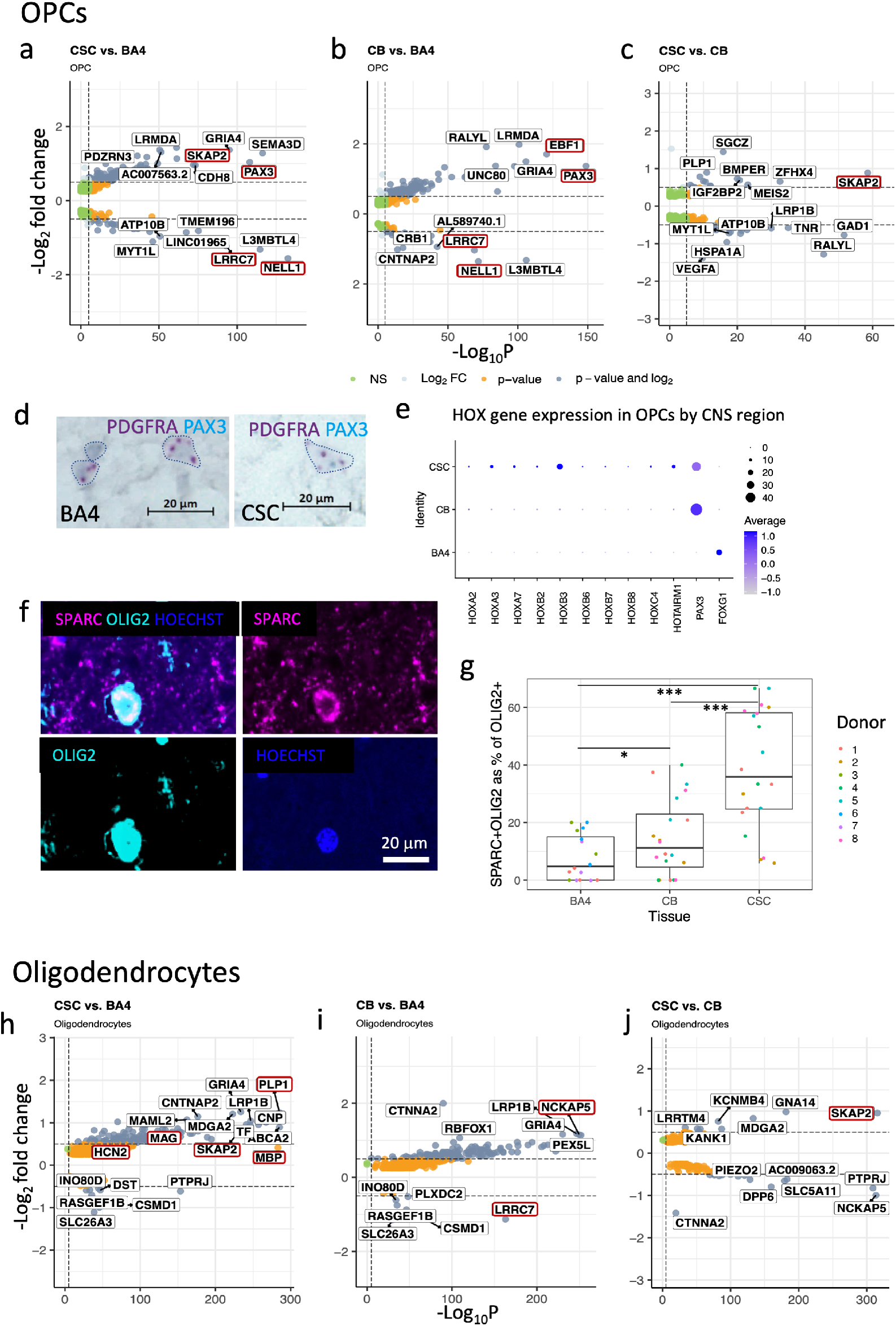
Regional oligodendroglial differences. a-c) Pairwise regional differentially expressed genes with region in OPCs. d) BaseScope duplex of *PAX3*-positive OPC in CSC but not in BA4. e) OPC *HOX* gene expression depending on CNS region in comparison with *PAX3* and the anterior marker *FOXG1*. f) Immunofluorescence of *SPARC*-positive *OLIG2*-positive oligodendrocyte (Oligo_F). g) Quantification of *SPARC*-positive Oligo_Fs N=5 (4 for BA4) in average counts per individual donor averaged across 4 fields of view. Visualised are the median, the 25th and 75th percentile (box) and minimum and maximum value (whiskers). Colours represent individual donors. h-j) Pairwise regional differentially expressed genes with region in oligodendrocytes.

Oligodendrocytes also vary with tissue region, notably with *SPARC*-expressing Oligo_F cells being significantly more abundant in the spinal cord compared to BA4 and CB (Figure 5a) confirmed by immunofluorescence co-labelling of *SPARC* and *OLIG2* on FFPE samples from different donors (Figure 5f,g, Table S4). The myelin sheaths around spinal cord axons are longer and thicker than those in the brain even on the same diameter axon^5^, and now this can be explained by the abundance of Oligo_F oligodendrocytes in the CSC, as these express more genes important for cholesterol biosynthesis and myelination (similarly to mice^60^), including *PLP1, CNP*, and *MAG* and of *HCN2* (known to control mouse myelin sheath length^33^) (Figure 2c–d, Figure 5h, Extended Data Figure 2c,Table S3). Differential gene expression analysis also showed that CSC OPCs and oligodendrocytes expressed more Src Kinase-Associated Phosphoprotein 2 gene (*SKAP2*), important for OPC mobility and myelin sheath formation in mouse spinal cord^61^ (Figure 5a,h). CB oligodendroglia showed increased differential expression of genes associated with synapse organisation and cellular adhesion, including NCK Associated Protein 5 (*NCKAP5*) and BA4 oligodendroglia expressed more Leucine-Rich RepeatContaining Protein 7 (*LRRC7*) (restricted to OPCs, COPs and Oligo_A) (Figure 5i,j).

Astrocytes also showed strong heterogeneity with WM tissue region: Clusters AS_5 and 6 were specific for BA4, clusters AS_4 and 7 were specific for CB and clusters AS_1, 2, 3, 9_ep, 11 and 12 were more abundant in CSC (Figure 4b,d) and so DEG analysis between regions reflected differences in these cluster marker genes such as increased *ADGRV1* (AS_5) in BA4 (Figure 6a–c, Table S5). In addition, CSC astrocytes expressed more Glial Fibrillary Acidic Protein (*GFAP*) as previously reported in mouse^62^ and more *SKAP2*, similarly to oligodendroglia (Figure 6a,c, Table S5).This astrocyte regional specificity may relate to the known variation in astrocyte density in different WM tracts and may help understand regional variation in human astrocytopathies^63^. For example, mutations in the *FAT3* gene are associated with Spinocerebellar Ataxia (SCA) and this is more highly expressed in CB astrocytes, (Figure 6a, Table S5), with increasing expression with age (Figure 7d).

**Figure 6.**
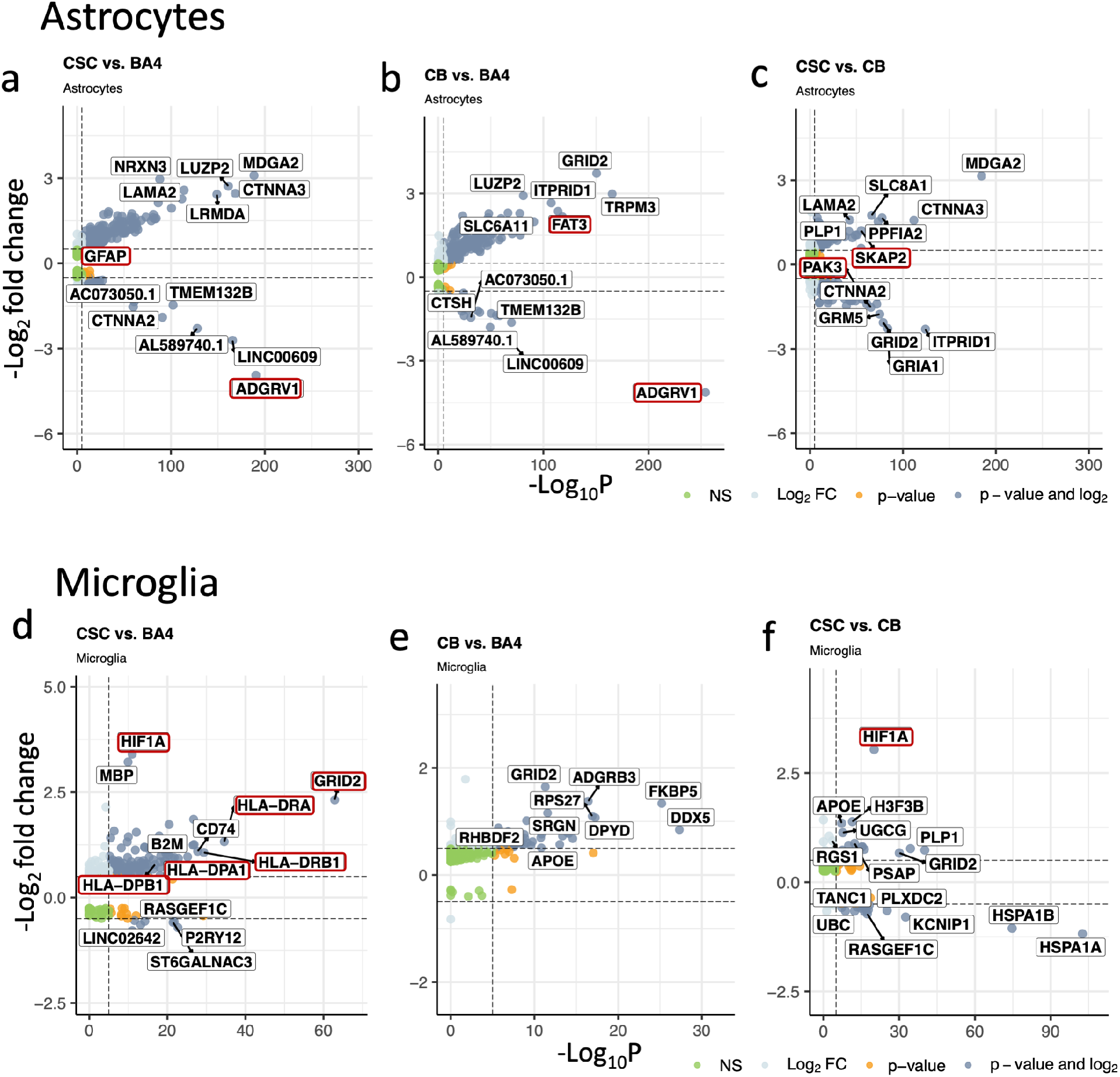
Regional astrocyte and microglial differences. Differential gene expression with CNS region in astrocytes (a-c) and microglia (d-f)

**Figure 7.**
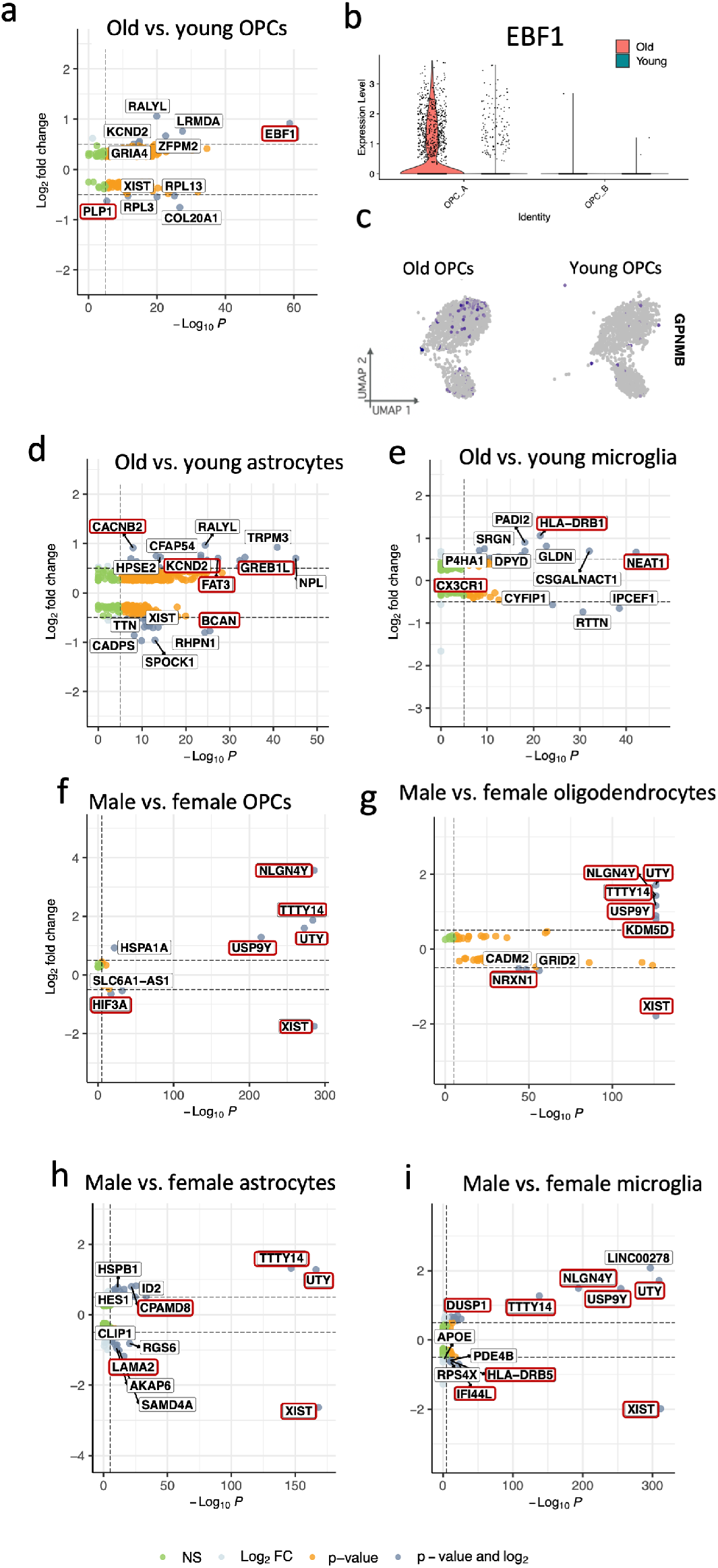
Age and sex effects in white matter glia. a) statistically significantly differentially expressed genes with age in OPCs include *EBF1* (b). c) Senescence marker *GPNMB* expression in OPCs of old and young donors. Differentially expressed genes with age in microglia (d) and astrocytes (e). Differentially expressed genes with sex in OPCs (f), oligodendrocytes (g), astrocytes (h) and microglia (i).

For microglia, the homeostatic Microglia_1 cluster was found equally in all three tissues, but activated Microglia_2 and Microglia_4 were mostly found in CSC followed by CB (Figure 4c,d). CSC microglia express more major histocompatibility genes and more *CD74* than in other regions, particularly BA4 (Figure 6d–f, Extended Data Figure 8), and perhaps both of these findings are as the blood spinal cord barrier is more permeable than the BBB resulting in an increased need for microglial responses to blood borne stressors. CSC and CB microglia also express more *GRID2* previously detected in brain microglia of AD donors^64^ and more *HIF1A* and *NEAT1* which may indicate a different response to hypoxic stress (Figure 6d–f, Table S5, Table S6, Table S7). Border associated macrophages (BAM) were mostly captured for CSC (Figure 4c,d) probably as meninges, including the perivascular space in which BAMs reside, were captured for this region.

### Age and sex differences are more subtle in healthy WM glia

We next tested for compositional differences with age and sex using Milo^57^ and using DEG analysis across the cell lineage specific dataset, then for each cellular subtype (each of the 60 clusters) separately, and identified genes of interest (Table S5, Table S6). Compositional differences were subtle (Extended Data Figure 9) but for age, DEG analysis showed that Early B-cell factor-1 (*EBF1*) was more highly expressed in OPC_A of older donors (Figure 7a,b). *EBF1*, although originally described in the haematopoietic system, is involved in Schwann cell myelination^65^, axonal pathfinding^66^ (such proteins are often similarly used for OPC migration) and specific polymorphisms of the gene associate with diagnosis of multiple sclerosis^67^. Older group spinal cord OPCs (OPC_A) also expressed less of the early myelin protein gene *PLP1* (Figure 7a), suggesting a decline in human OPC function with age, as in aged rat OPCs^10^. This functional rat OPC decline with ageing was related to increased senescence markers^10^ but in our dataset there was no statistical difference in the expression of known markers of cellular senescence between age groups in OPCs (or indeed in any cell lineage) including P16 (*CDKN2A*), P21 (*CDKN1A*), P53 (*TP53*), senescence-associated beta-galactosidase (*GLB1*) and Glycoprotein Nmb (*GPNMB*). However, expression of *GPNMB* shown on a feature plot suggests an increase with age in OPCs (Figure 7c). Differentially expressed genes with older age in astrocytes include *CACNB2* and *GREB1L*, important for calcium signalling and the potassium voltage-gated channel protein gene *KCND2* (Figure 7d). Astrocytes of younger donors express more *BCAN* which is important for experience-dependent neuroplasticity and normal cognitive function^68^ (Figure 7d). Previous literature suggests that microglia express more activation genes with increased age^12^, and our data confirms this, for example, *HLA-DRB1* and *HLA-C* are more highly expressed in old microglia (Figure 7e) and markers of homeostasis such *CX3CR1* decline with age (Table S5). However, this relatively subtle difference in activation may be bigger in vivo as snRNAseq may underestimate the expression levels of microglia activation genes in comparison to scRNAseq^69^. Differences in gene expression with sex were largely restricted to the expression of gonosomal genes, such as *XIST* in female donors and Y-chromosomal genes in glia of male donors: *NLGN4Y, TTTY14, USP9Y, KDM5D, ZFY* and *UTY* (Figure 7f–i). Albeit these are not surprising, they may still be important, as studies have shown a potential effect of gonosomal genes on neuropathologies: when Y-linked genes were repressed in a mouse model of Alzheimer’s disease, more mitochondrial gene dysfunction and neuroinflammation was observed^15^. However, in a study on AD patients and controls, very few differentially expressed genes between male and female microglia other than gonosomal genes were found^70^. Other differentially expressed genes with sex include more *HIF3A* in female OPCs (related to oxidative stress) (Figure 7f), more *NRXN1* in female oligodendrocytes (Figure 7g), more *LAMA2* in female astrocytes (related to the BBB) (Figure 7h), and more pro-inflammatory genes such as *HLA-DRB5* and *IFI44L* in female microglia (Figure 7i), with more *CPAMD8* (associated with late-onset Alzheimer’s disease^71^) in male astrocytes, (Figure 7h), and higher expression of *DUSP1* which modulates microglia towards a homeostatic phenotype^72^ in male microglia (Figure 7i). These changes suggest sex differences in the link between inflammation and ageing, and may explain some sex dimorphism in neurodegenerative disease susceptibility.

Therefore, in summary, we show that in normal adult human WM glia, there is marked variation in transcriptional signatures of all broad cell groups with region, finding both regionspecific, region-selective and human-specific cell subtypes, with much less variation with age and sex.

## Discussion

We predicted that there would be variation of glial transcriptional signatures as a read-out of different function across CNS regions, older age and between sexes, and this might help explain the influence of these factors on susceptibility to neurological disease. We found marked regional differences between the spinal cord and the brain, so that we can no longer assume parity of physiological, pathological or therapeutic response between the two.

The clear difference between posterior (CSC and CB) and anterior OPCs (BA4), and the continued expression of *PAX3* and *HOX* genes even in adulthood in these spinal cord OPCs, strongly suggests that these are derived similarly to in mouse where *Pax3* identifies the third wave of OPCs from a dorsal origin^34^. We speculate that these OPCs differentiate into Oligo_F oligodendrocytes which are selective for the CSC and seem only present in humans. This Oligo_F cluster is notably different from the other mature oligodendrocyte clusters, with higher expression of myelin gene mRNAs and *HCN2*, in line with our knowledge that spinal cord oligodendrocytes produce more myelin, in the form of longer and thicker myelin sheaths per axon diameter, than brain oligodendrocytes^5, 33^. Remyelination in the spinal cord after damage in MS appears less successful than in the brain^73^, which increases our interest in carrying out snRNAseq analysis from spinal cord WM in donors with MS – are Oligo_F cells selectively lost and/or replaced by oligodendrocytes with less (re)myelination capacity? Or are myelin sheaths difficult to replace in the spinal cord as they need to be thicker and longer? Other differences in the CSC may relate to the more ‘open’ blood spinal cord barrier compared to the BBB therefore likely exposing CSC cells to more blood-derived inflammatory cells/factors which may explain the increased expression in immune-related genes in Oligo_F, but also the more activated state of CSC microglia, and alterations in CSC astrocytes. These differences may clearly be of relevance to diseases which either selectively/predominantly affect the spinal cord, and for best targeting of therapeutics.

Age differences were more subtle and we hypothesise that we will see bigger effects with a bigger age disparity, perhaps if we included children/adolescents as our ‘young’ group. However, some interesting genes such *EBF1* have emerged which may be useful as a marker of aged adult OPCs, which we know (in rat) are poorer at responding to damage and initiating remyelination than young ones^74^. Some changes with age or sex may also affect lowly expressed genes, which may be hard to detect by snRNA-seq^69^. Cell sorting strategies on larger cohorts based on markers identified in the present study with subsequent bulk RNAseq could highlight such differences in lowly expressed genes. The possibility also remains that some physiologically relevant age and sex-related differences may not lie in the transcriptome and be, instead, post-transcriptional and/or metabolic. Finally, determining whether the subtypes we discovered are associated with local microenvironment or specific cell-cell interactions will require more spatially-resolved techniques.

Our results show that human post-mortem tissue is an invaluable source of information not obtainable otherwise because of human-specificity of some cellular phenotypes (such as Oligo_F), and difficulties in establishing faithful human in vitro models. However, by definition, human postmortem tissue only presents a snapshot of transcriptional activity at a single moment in time. Our attempts to infer transcriptional changes over time (trajectory inference) for the oligodendroglial lineage, where OPCs and mature oligodendrocytes co-exist in the adult CNS, showed no clear trajectory between or within these cells. This may be due to low levels of oligodendroglial turnover with few intermediates in the non-diseased, adult human CNS, supported by a lack of markers of cell division in OPCs, and consistent with previous radioisotope studies^75^. Also, oligodendrocyte subtypes may either have fixed transcriptional profiles in adulthood, or may dynamically alter their transcriptional profiles in multidirectional ways, obscuring simple trajectories. We know that oligodendroglial transcriptional change is possible in response to disease, by the appearance of new oligodendrocyte clusters found in MS donors, but we do not yet know whether these are generated from other oligodendrocytes transitioning between clusters or whether they are newly-formed from OPCs in response to damage. The presence of more intermediate oligodendroglia in the MS dataset^25^ compared to our current dataset suggests an increase in the trajectory from OPCs to oligodendrocytes, as classically described^76^. However, there is also increased evidence for and interest in the ability of mature oligodendrocytes to extend processes and form new myelin in response to demyelination, likely needing additional transcriptional changes^75, 77, 78^. These, not mutually exclusive, options will only be solved by use of reporter genes for identified clusters in human cell models.

Our findings in normal post-mortem WM of marked regional cellular and gene expression differences are critical for the understanding of diseases which are regionspecific/selective and to generate more effective therapeutic strategies. Our results would predict that in MS, for example, different pro-remyelinating therapies may be effective after spinal cord demyelination compared to brain. Our analysis will allow appropriate comparisons with future diseased human CNS cohorts and we provide these data as an open resource for others in line with the Human Cell Atlas Project, and also in a Shiny app for easy browsing.

## Methods

### Human donor tissue

Adult post-mortem unfixed fresh-frozen tissue and formalin-fixed paraffin-embedded (FFPE) tissue were obtained from the MRC Sudden Death Brain Bank in Edinburgh with full ethical approval (16/ES/0084) from a total of 20 Caucasian and British donors. Fresh-frozen samples of two different age groups, “young adults” (30-45 y, 5 males and 5 females) and “old adults” (60-75 y, 5 males and 5 females), were used and selected so that following three tissue regions were present for each donor: motor cortex (Brodmann area 4, BA4), cerebellum (CB) and cervical spinal cord (CSC) (Extended Data Table 1). FFPE sections (4 µm) of the same regions were from a separate group of individuals of the same age groups and both sexes (Extended Data Table 1). These samples had been verified to have no neuropathology by a consultant neuropathologist (Professor Colin Smith).

### Nuclei isolation

For each sample, 20 tissue cryosections at 20 µm thickness were macro-dissected and the WM (identified using Luxol Fast Blue staining) was used for nuclei isolation. For BA4 we selected sub-cortical white matter, for CB the stem of the arbor vitae for CSC the big dorsal white matter tracts: fasciculus cuneatus and fasciculus gracilis. The nuclei PURE isolation kit (Sigma, NUC201-1KT) was used with following the manufacturer’s instructions with slight alternations, explained in Supplementary File 1. Nuclei were stained with Trypan blue (Bio-Rad Laboratories, 1450013) and their concentration was measured using an automated cell counter (Biorad TC20) and standardised to 1,000,000 nuclei per ml.

### 10X loading and library preparation

Samples were randomly allocated to 10X Genomics Chromium single cell 3’ chips. The 10X Genomics v3 beads, Gem kits and library kits were used according to the manufacturer’s instructions. Library quality was assessed using a Bioanalyzer (PerkinElmer LAS (UK) Ltd). See Supplementary File 1 for more information.

### Next Generation Sequencing

Nine to ten cDNA libraries were randomly allocated to seven sequencing pools and added at an equimolar concentration of 5 nmol. Sequencing was performed by Edinburgh Genomics using a Novaseq 6000 System and a S2 flow cell. Each pool was sequenced on two different lanes to reduce batch effects. We aimed for a sequencing depth of 70,000 reads per nucleus.

### Genome alignment and raw matrix generation

The transcriptome was demultiplexed and aligned with the human reference genome GRCh38 using 10X Genomics Cell-ranger 3.0.2.

To retrieve feature-count matrices for exonic and intronic reads separately Velocyto (0.17.16) was used with a repeat mask for the human reference genome GRCh38 which was downloaded from https://genome.ucsc.edu/index.html.

### Data Quality control

Genes with less than 200 copies across both the spliced and unspliced matrices were removed from the analysis. The nucleus quality control was conducted on the spliced and unspliced matrices separately using UMI counts, gene counts and percentage of mitochondrial RNA. Table S1 visualises used thresholds and how many nuclei were filtered at any step and in total. Library sizes did not correlate with RNA integrity values (Extended Data Figure 1c).

### Normalisation

Scran was used to normalise the combined matrix of spliced and unspliced reads for each tissue separately and to transform data onto a logarithmic scale.

### Dimensional reduction and clustering

Using Seurat^39^, the 2000 most variable genes were selected, all genes were scaled and a principal component analysis was performed. The first 25 principal components were used for constructing a Shared Nearest Neighbour (SNN) graph (k = 20) and as input for a Uniform Manifold Approximation and Projection (UMAP). This process was repeated after sample and cluster quality control steps. Louvain clustering was performed at a resolution of 0.8 (0.5 and 1.3 were also tested). The clustering behaviour was not influenced by the allocation of a sample to an individual chromium chip or sequencing batch (Extended Data Figure 1g) but samples varied in quality. A combination of following statistics was used to filter for good quality samples: mean UMI count > 490, proportion of spliced vs. unspliced genes < 75 %. This resulted in ten samples being removed from downstream analyses (4 BA4, 5 CB, 1 CSC; 6 young, 4 old; 5 male, 5 female) (Extended Data Figure 1d–f). We considered following criteria to manually curate clusters: Clusters had to contain nuclei of at least 8 donors, while donors were only considered if they contributed at least 2 % to the total cluster size.

### Cell type annotation and analysis of subsetted cell lineage datasets

Canonical marker genes were used to annotate cell types as described in the main text and to select oligodendroglia, astrocytes, microglia and macrophages for separate downstream analysis. The 2000 most variable genes were selected, all genes were scaled and a PCA was performed. The first 10 principal components were used for constructing a SNN graph with 20 knearest neighbours and as input for a UMAP. Our clustering strategy involved repeated clustering at different resolutions from obvious under- to clear over-clustering and was considered in combination with differential gene expression analyses (see below) and measures of cluster stability at all those resolutions as described in Supplementary File 1. If functional differences could be recognised based on the expression of marker genes, they were considered in the annotation (e.g. excitatory vs. inhibitory neurons).

### Differential gene expression analysis

Differential gene expression analyses for the identification of cluster marker genes and genes that are enriched in tissue, age and sex groups were performed using MAST^79^ within Seurat (using test.use = “MAST”) filtering genes for those that had a minimum positive log2-fold change of 0.25 and were expressed by at least 25 % of cells within the cluster/group of interest. All genes that were expressed in less than 60 % of nuclei outside the cluster/group of interest were considered for the visual screening of marker genes and as gene ontology input if they had a minimum log2-Fold change of 0.7 and 0.4 respectively.

### Compositional analysis

We used the R library miloR^57^ for differential abundance testing as specified in Supplementary File 1 to investigate if the number of nuclei per cluster correlated with tissue, age or sex group.

### Label transfer and integration with other datasets

Seurat’s CCA label transfer and integration was applied to our oligodendroglia dataset and a published human^25^ or three mouse datasets^2, 3, 40^. For the label transfer and the integration, the first 30 and 20 dimensions respectively were used to specify the neighbour search space. The integrated data was re-scaled and dimensionally reduced using the first 20 principal components as input for a UMAP.

We created a correlation matrix between predicted and current labels by counting each occasion of a co-labelling and normalising counts for cluster size and largest cluster to bring the results on a scale from zero to one. The R library corrplot was used with the argument is.corr = FALSE to visualise cluster label correlations.

### Trajectory analysis

Dynverse^35^ was used for the selection of suitable trajectory inference methods for the complete oligodendroglia dataset and the oligodendroglia dataset divided by tissue. Slingshot^37^, PAGA and PAGA tree, Angle and Scorpius were the top ranking methods for our dataset based on Dynverse and they were all tested within the Dynverse framework, setting PDGFRA expressing OPC nuclei as start point for the trajectories. Additionally, slingshot^37^ and Monocle^36^ were run outside of Dynverse^35^ first separately for each tissue (BA4, CB, CSC), then for the tissues combined. In Monocle, the starting point was first not determined, but because we know that in vitro OPCs can generate oligodendrocytes, we set the start point to the OPC clusters. Lastly, we used scVelo^38^ within R using velociraptor^80^ on spliced and unspliced counts that were subsetted for nuclei that were retained in the final dataset. We superimposed UMAP projections onto the new dataset that included RNA velocity^81^ calculations.

### Gene Ontology

Cluster profiler^82^ was used on all significantly differentially expressed genes (adjusted p-value < 0.05) with a log2-Fold change of at least 0.4 (which corresponds to at least approximately 30 % difference) to identify significantly enriched biological processed in clusters, and tissue, sex and age groups as described in Supplementary File 1.

### Computational analysis systems

Earlier steps of the analysis (genome alignment) and computational intensive analyses (such as Monocle^36^ on oligodendroglia of all tissues) were performed on the University of Edinburgh’s Linux compute cluster. For later steps R (version 4.1.1.) and RStudio (version 1.3.1056) were used on a Macbook Pro (64-bit) with macOS Big Sur 11.6.1. More information on library versions used can be found in the github repositories.

### Data availability

All data necessary to reproduce the results of the present paper will be available after publication. R code (that also specifies R library versions), a shiny app visualising the complete and all cell lineage datasets and supplementary tables showing differential gene expression and gene ontology analyses results will be available as well.

### Immunofluorescence

FFPE tissue sections were de-waxed and re-hydrated in 2 x 10 min in Xylene, 2 x 2 min in 100 % ethanol, then 2 min in each 95 %, 80 % and 70 % of ethanol. Antigen retrieval was performed by microwaving the samples for 15 minutes in 0.01 M citrate buffer (pH = 6, Vector Laboratories, H-3300) in Millipore water. Sudan Black Autofluorescence Eliminator Reagent (Merck Millipore, 2160) was added to each slide for 30 s – 1 min which were then washed twice for 5 min in TBS with 0.001 % Triton. Slides were incubated in 3 % H2O2 for 10 min and washed. Serum block (TBS + 0.5 % Triton + 10 % horse serum) was added for 1 h at room temperature. Antibodies were diluted in serum block and added to slides prior to an overnight incubation at 4°C. On the next day, slides were washed in TBS + 0.001 % Triton (2 x 5 min) and incubated with an ImmPress HRP secondary antibody (Vector Laboratories, MP-7401-15) for 1 h. After washing the slides, OPAL 570 tyramide dye (Akoya Biosciences, SKU FP1488001KT) at a dilution of 1:500 was added for 10 minutes. After washing them, the slides were boiled in 0.01 M citrate buffer (pH = 6) for 2.5 minutes and then left in the hot buffer for 20 minutes. Slides were washed (2 x 5 min) and incubated overnight at 4°C with the second primary antibody diluted in serum block. On the third day, slides were washed (2 x 5 min), incubated with appropriate Vector ImmPress HRP secondary antibody (Vector Laboratories, MP-7405) for one hour at room temperature, washed again (2 x 5 min) and incubated with OPAL 650 Tyramide (Akoya Biosciences, FP1496001KT, 1:500) for 10 minutes. After washing the slides (2 x 5 min) they were counterstained with Hoechst (10 min), washed (2 x 5 min) and mounted.

Both, the SPARC (Abcam, ab225716) and the Olig2 (R&D Systems, AF2418) antibodies were used at a 1:100 dilution.

### BaseScope duplex

We used a BaseScope duplex detection kit (ACD Bioscience, 323800) on FFPE tissue according the manufacturer’s instruction with following optimised times: Target retrieval for 20 min, AMP7 and AMP11 incubation for 1h each. We used following probes: *PDGFRA* and *PAX3*.

### Imaging and analysis of imaging data

A Vetra Polaris slide scanner (Perkin Elmer) was used to image the IF at 20x objective. Four fields of view within WM were used for each sample to manually quantify tissue stainings in QuPath^83^. BaseScope duplex stainings were imaged using an inverted Widefield Observer (Zeiss). The SPARC/OLIG2 validation was analysed using Poisson models with SPARC+OLIG2+ counts as response variable. Following predictors were tested: Tissue region, total OLIG2+ count, age (as factor with the two levels “young” and “old” and as covariate) and sex as fixed effects and donor ID and field of view tested as random effects. All non-significant predictors were removed from the model (based on F-test for fixed effects and covariates and Log-likelihood ratio test for random effects). The final model was a generalised linear Poisson model with Tissue and total OLIG2+ count as predictors.

## Acknowledgements

This project was funded by a Medical research Council grant as part of the Human Cell Atlas Project HCA, by grant number CZF2019-002427 from the Chan Zuckerberg Foundation by the MS Society UK Edinburgh Research Centre and from a British Heart Foundation pilot grant. This work has made use of the resources provided by the Edinburgh Compute and Data Facility (ECDF) (http://www.ecdf.ed.ac.uk/).

We thank the following people and facilities at the University of Edinburgh for their help: The MRC brain bank for providing tissue, Pamela Brown from the Biomolecular Core facility for Bioanalyzer measurements, SuRF Histology for cutting paraffin samples, and Matthieu Vermeren and Justyna Cholewa-Waclaw from the CRM imaging facility for their continued support with imaging tissue. Next Generation Sequencing of cDNA was carried out by Edinburgh Genomics at the University of Edinburgh. Some figures were created with BioRender.com. We also thank Alex Lederer and Alan O’Callaghan for discussions about the bioinformatics analysis and their advice.

## Author contributions statement

LAS and SJ conducted the snRNAseq experiment and LAS analysed the data with support from NBC and input from NLK, SB, AMK, DVB, MK, FBP and ZM. LAS did the IF and BaseScope validation. NH provided access to a 10X chromium controller and reagents and provided advice. AW, GCB, GL, CAV secured funding and supervised the project. LAS and AW wrote the first manuscript version and all authors contributed with their comments to the submitted version.

---

**Extended Table 1.**
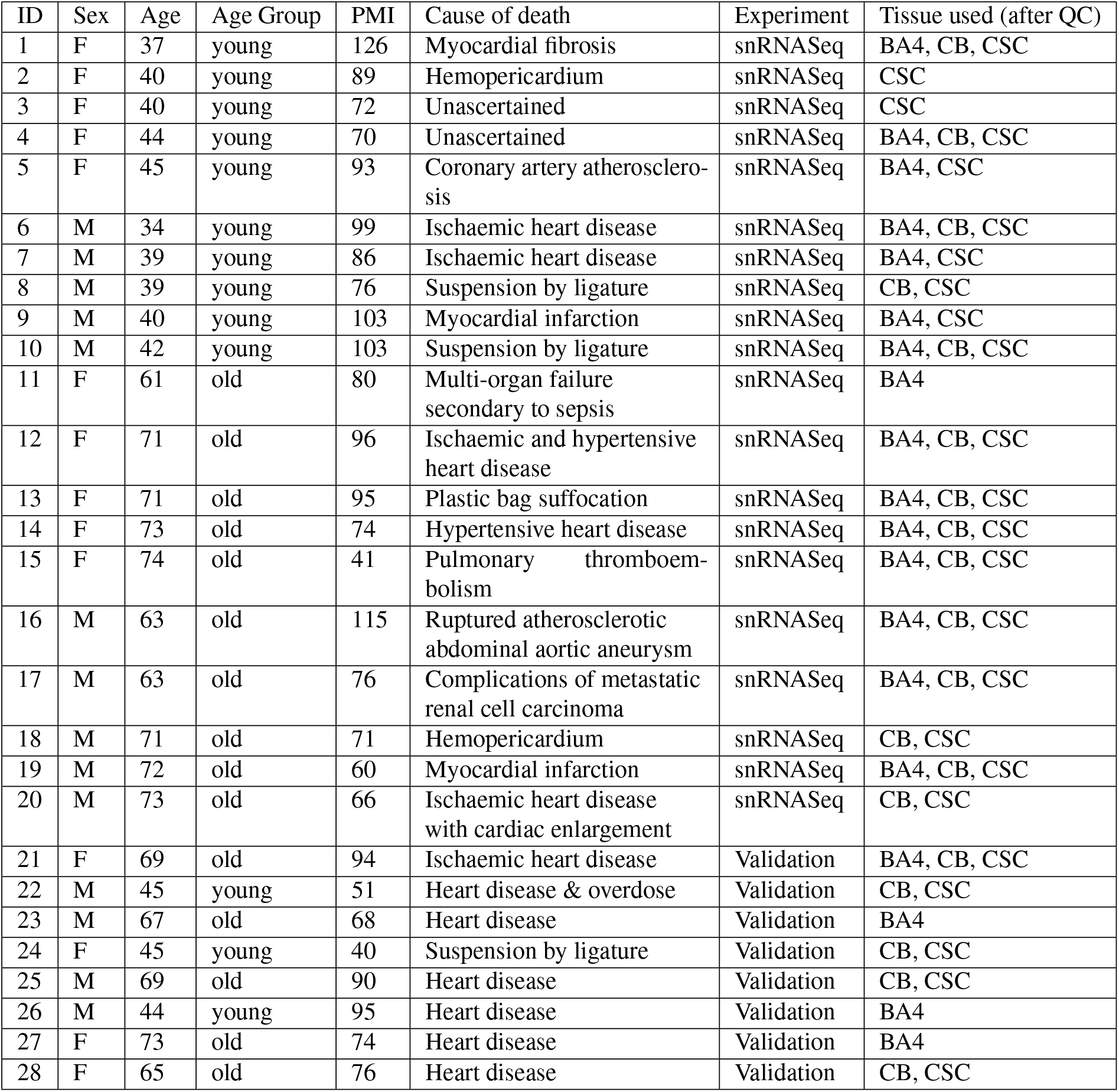
Donor information for snRNAseq experiment. PMI: post mortem interval in hours.

**Table S 1.** Nuclei quality control on spliced and unspliced matrices separately. *Total number of nuclei filtered out during the nuclei quality control step; Some nuclei were filtered out during quality control steps for both the spliced and unspliced matrix and therefore this number does not represent the sum of the nuclei in the previous column. Note that mitochondrial genes are only present in the spliced matrix. In the combined matrix, the majority of samples had a mitochondrial percentage of less than 4%.

**Extended Table 1.** Donor information for snRNAseq experiment. PMI: post mortem interval in hours.
| Matrix | Filter level | Threshold | Number of Filtered nuclei | Total nuclei within matrix | TOTAL |
| --- | --- | --- | --- | --- | --- |
| spliced | Low UMI count | 100 | 95 | 16927 | 29028* |
|  | Large UMI count | 10156 | 1695 |  |  |
|  | Low gene count | 33 | 3429 |  |  |
|  | Large gene count | 6791 | 225 |  |  |
|  | High mitochondrial gene percentage | 23 | 15306 |  |  |
| unspliced | Low UMI count | 100 | 21749 | 21774 |  |
|  | Large UMI count | 30000 | 25 |  |  |
|  | Low gene count | 10 | 3912 |  |  |
|  | Large gene count | 8000 | 0 |  |  |

**Table S 4.** Statistical analysis output of Oligo_F validation using Poisson model. SE: Standard Error. CB: cerebellum, CSC: cervical spinal cord.

|  | Estimate | SE | z-value | p-value |
| --- | --- | --- | --- | --- |
| Intercept | -0.097 | 0.273 | -0.354 | 0.723 |
| Tissue: CB | 0.5 | 0.249 | 2.008 | 0.045 |
| Tissue: CSC | 1.483 | 0.226 | 6.557 | <0.0001 |
| total Olig2+ | 0.022 | 0.006 | 3.759 | <0.001 |

**Extended Figure 1.**
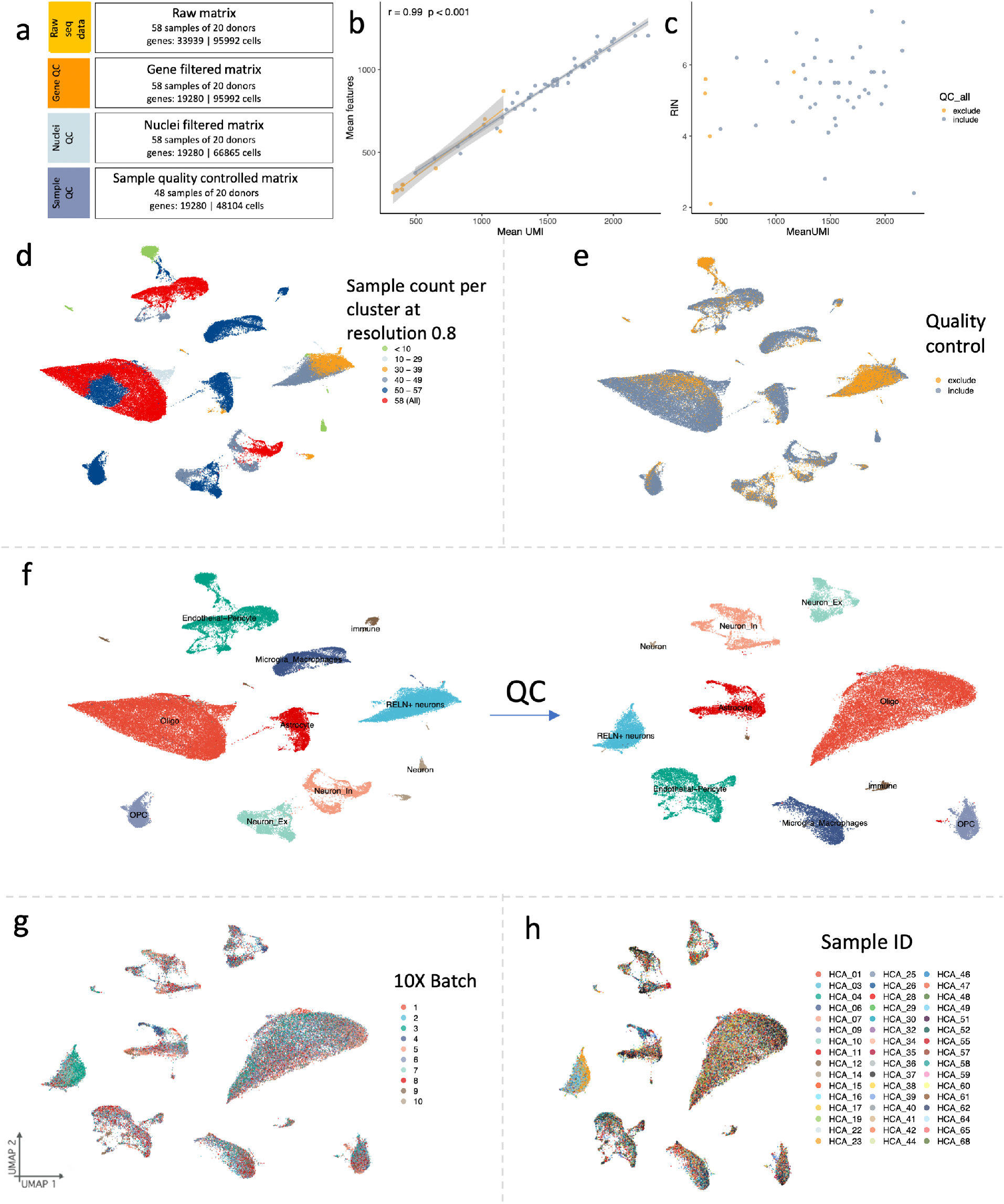
Quality control. a) Effects of quality control filtering on data size. b) Correlation of gene and unique molecular identifier(UMI) count within cells. c) No correlation of RNA integrity value (RIN) with library size (mean UMI count). Before cluster quality control, some clusters were due to few samples (d). e) Cells that were included/excluded in downstream analysis. f) Complete dataset before and after cluster quality control and re-clustering. g) No 10X batch effect was observed and h) clustering was not determined by individual samples after quality control

**Extended Figure 2.**
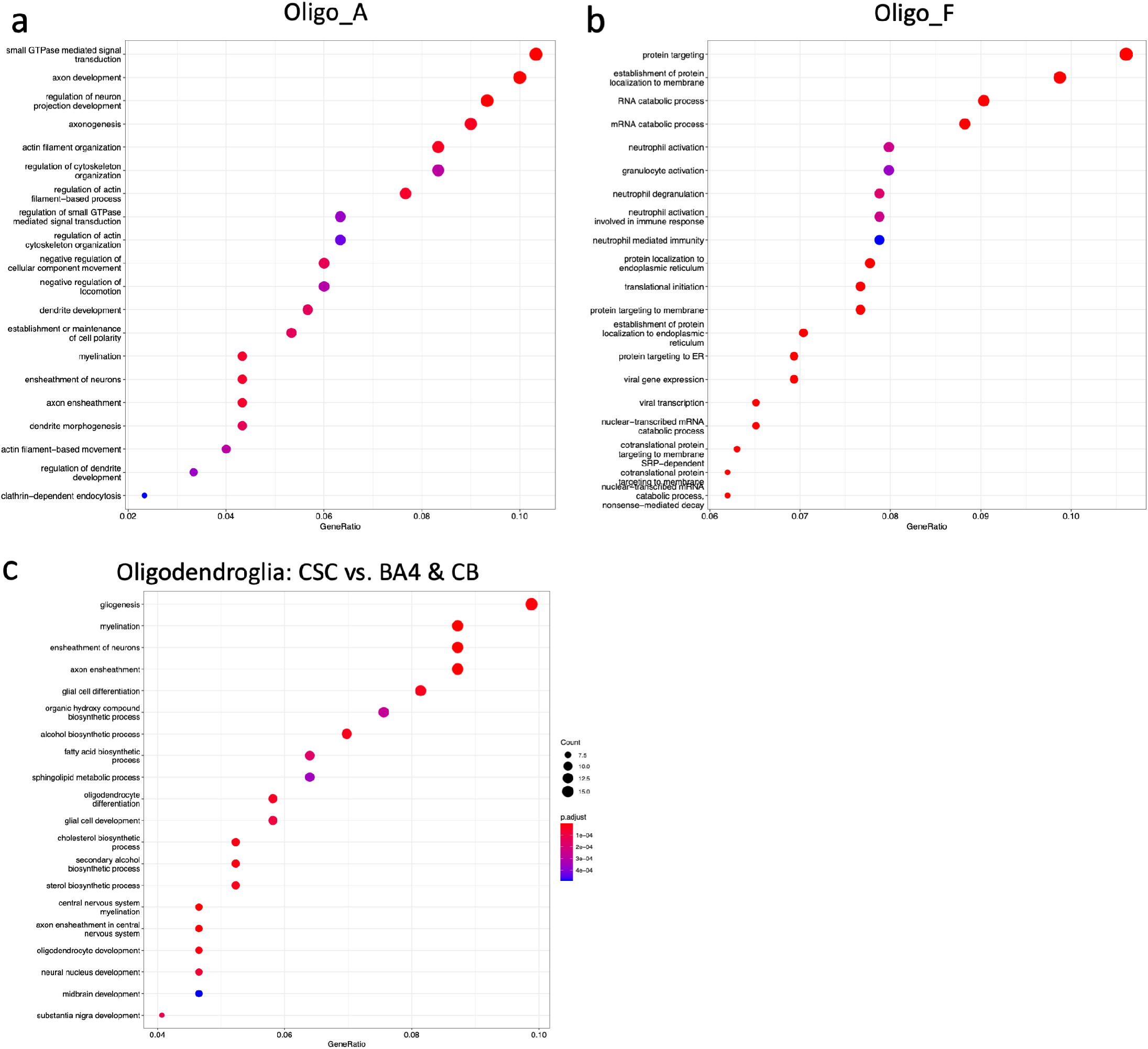
Oligodendroglia gene ontology terms associated with specific clusters a) Oligo_A and b) Oligo_F.

**Extended Figure 3.**
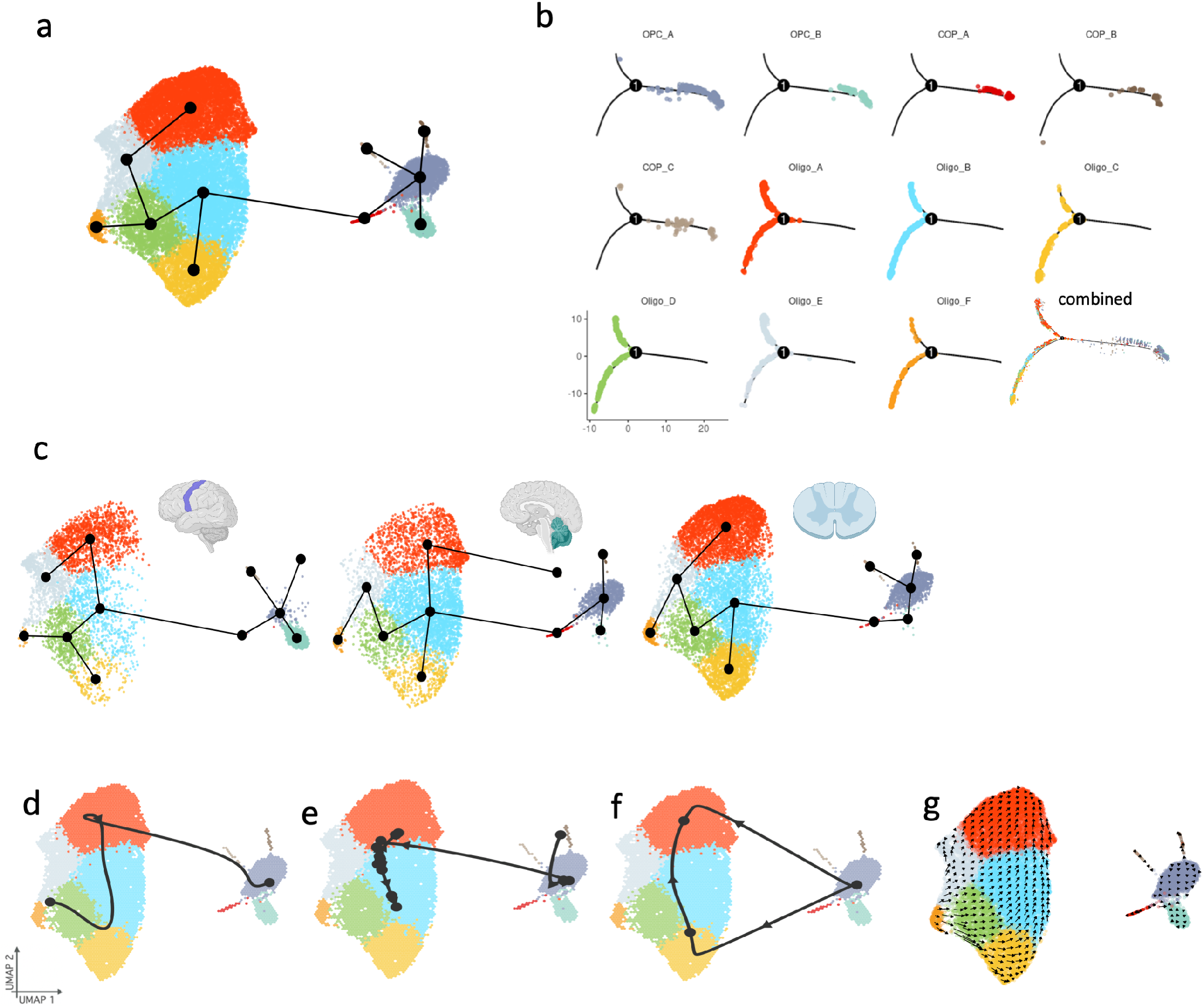
Trajectory inference methods such as Slingshot (a, c) Monocle (b), Scorpius (d), PAGA-tree (e), Angle (f) and scVelo (g) do not agree on a likely trajectory. All trajectory inference methods are shown for the complete dataset and Slingshot also individually by CNS tissue (c).

**Extended Figure 4.**
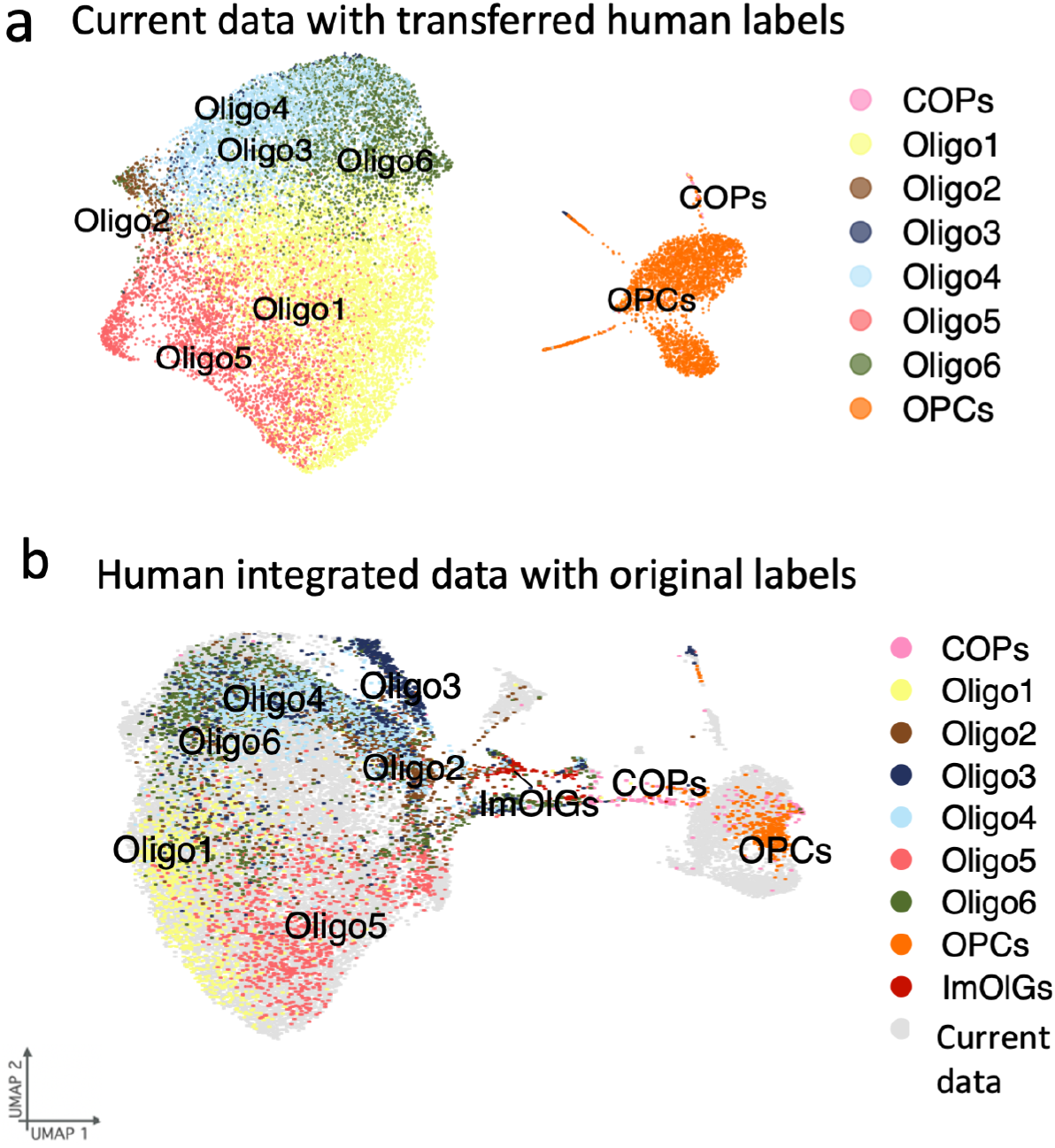
Correspondence of current dataset with previous human dataset25. a) Current data with transferred labels from ref.25. b) CCA integrated dataset coloured by previous labels.

**Extended Figure 5.**
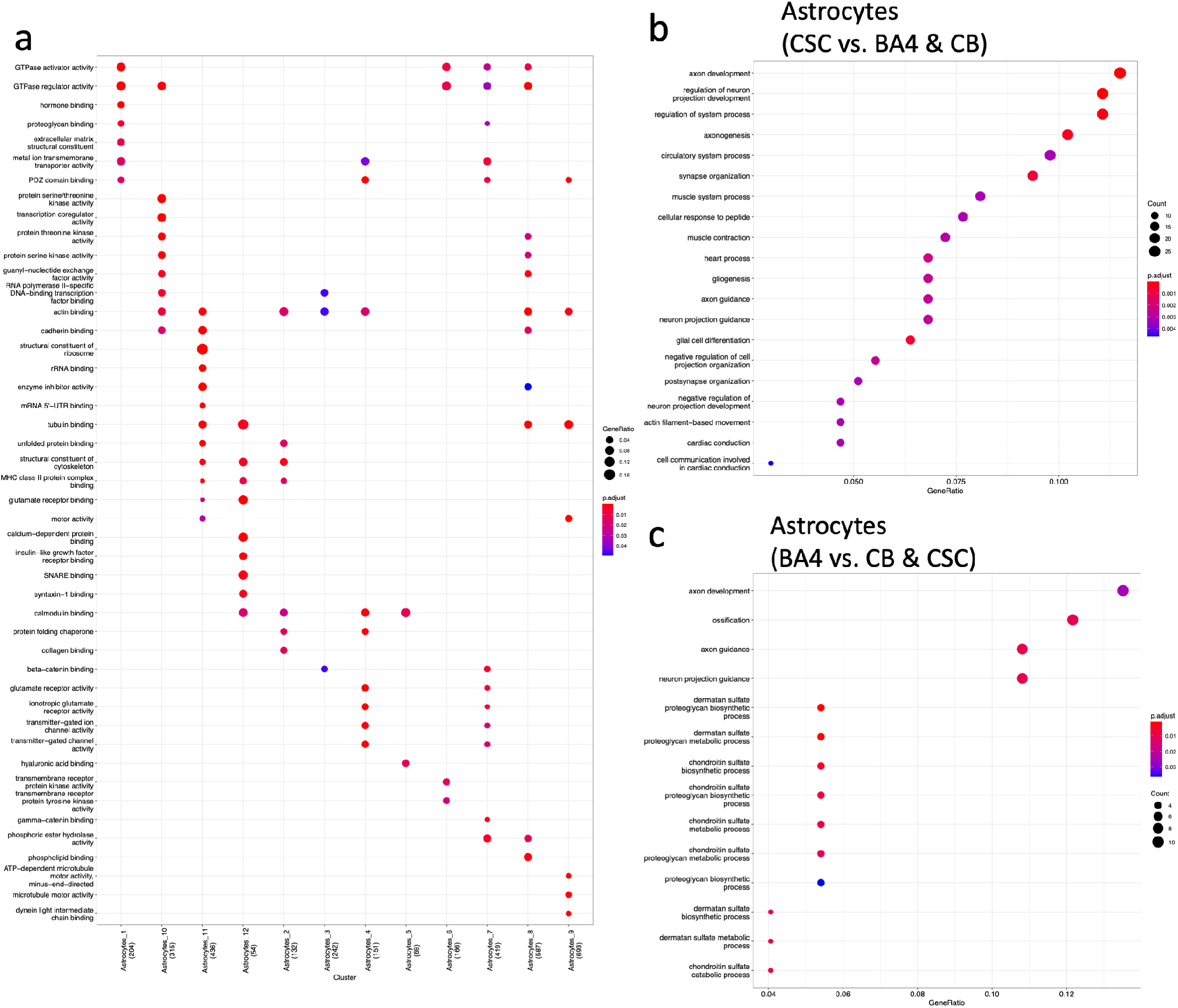
Astrocyte Gene ontology with a) cluster and b-c) CNS region.

**Extended Figure 6.**
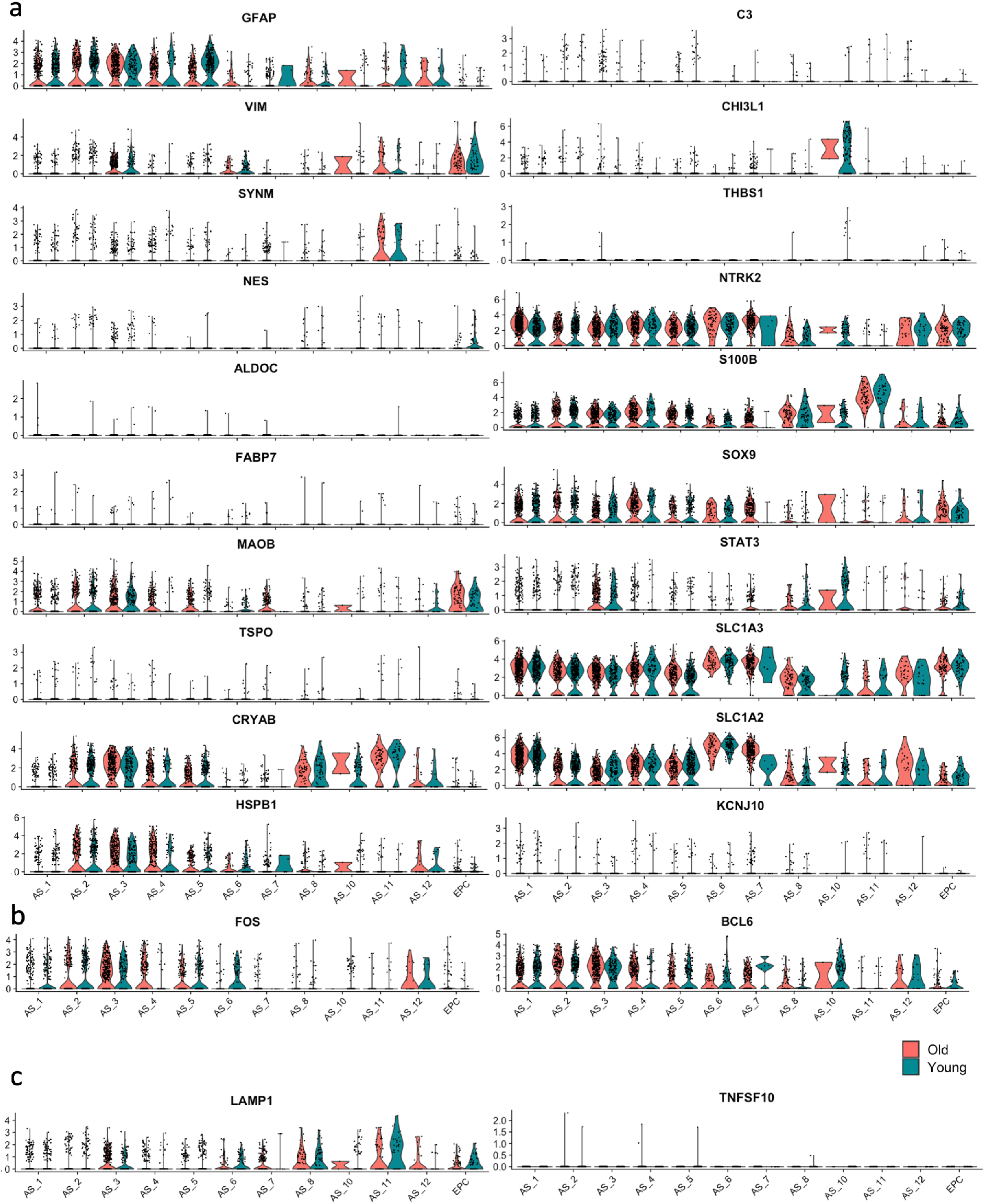
Activation state markers for astrocytes. a) Genes that are associated with a more activated state51. b) Markers that were upregulated at the borders of multiple sclerosis white matter lesions26 and c) genes that limit inflammation in the CNS and indicate a more inactivated state84

**Extended Figure 7.**
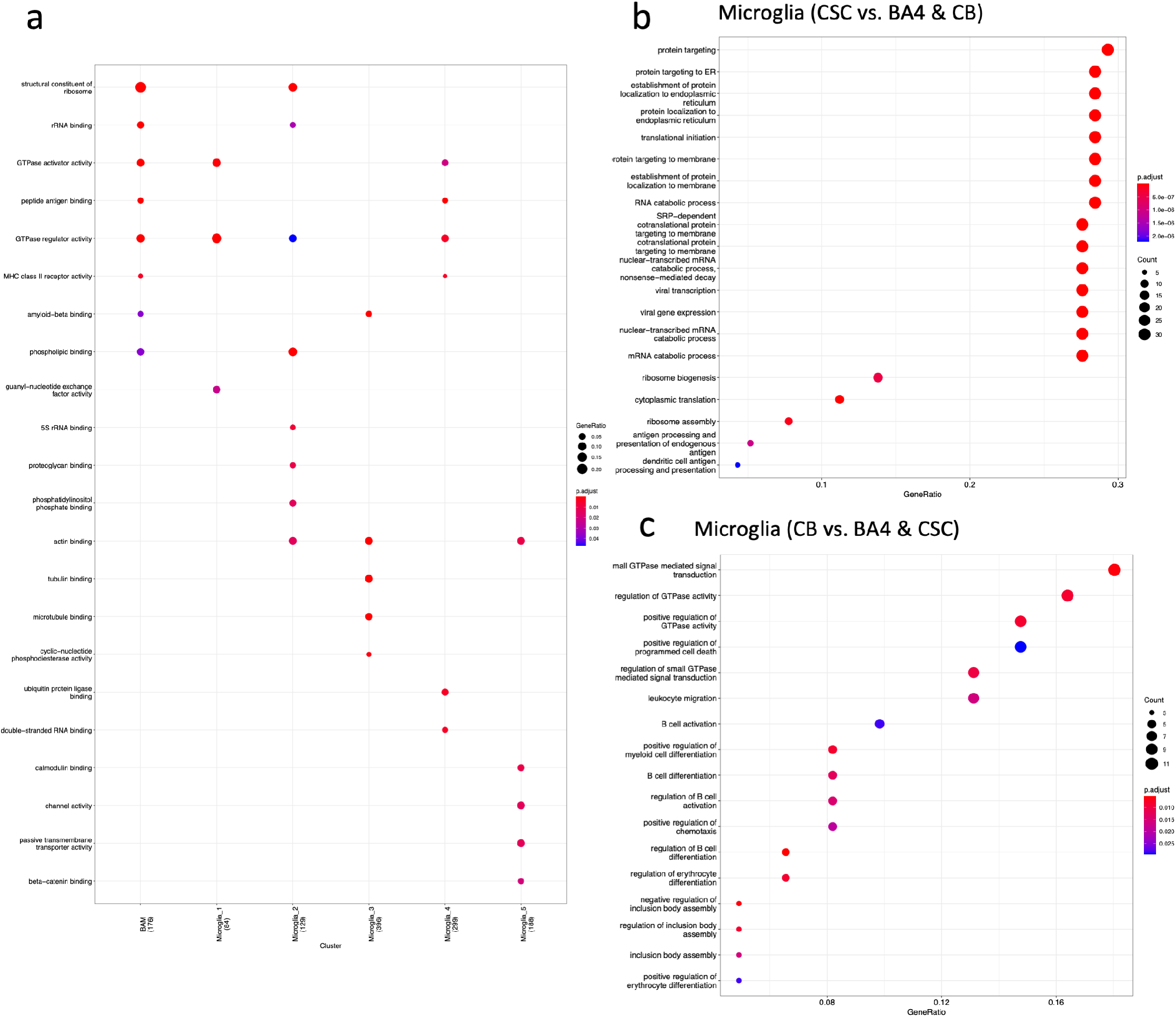
Microglia gene ontology with a) cluster and b-c) CNS region.

**Extended Figure 8.**
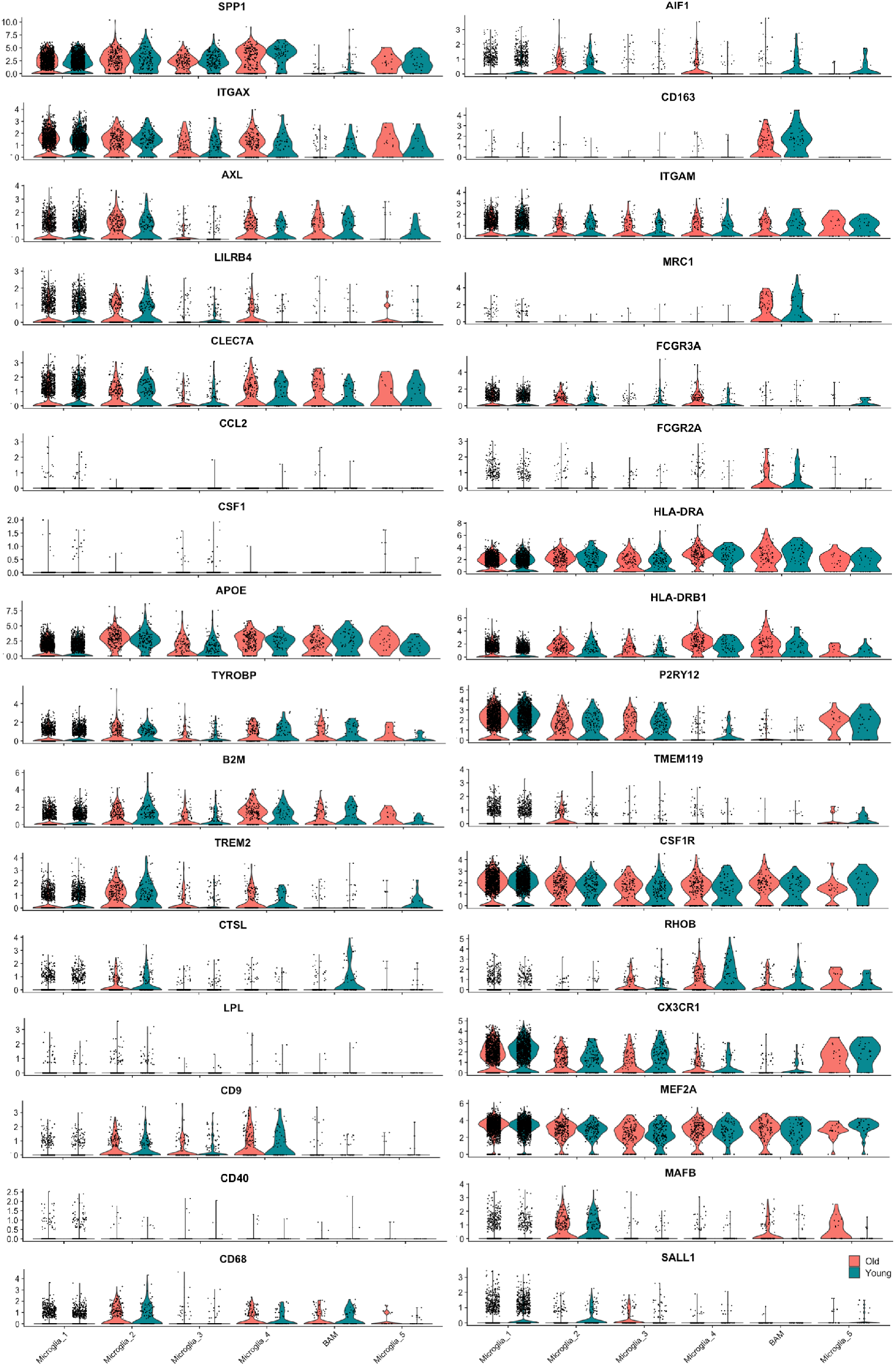
Microglia gene expression of markers associated with microglia activation and homeostasis.

**Extended Figure 9.**
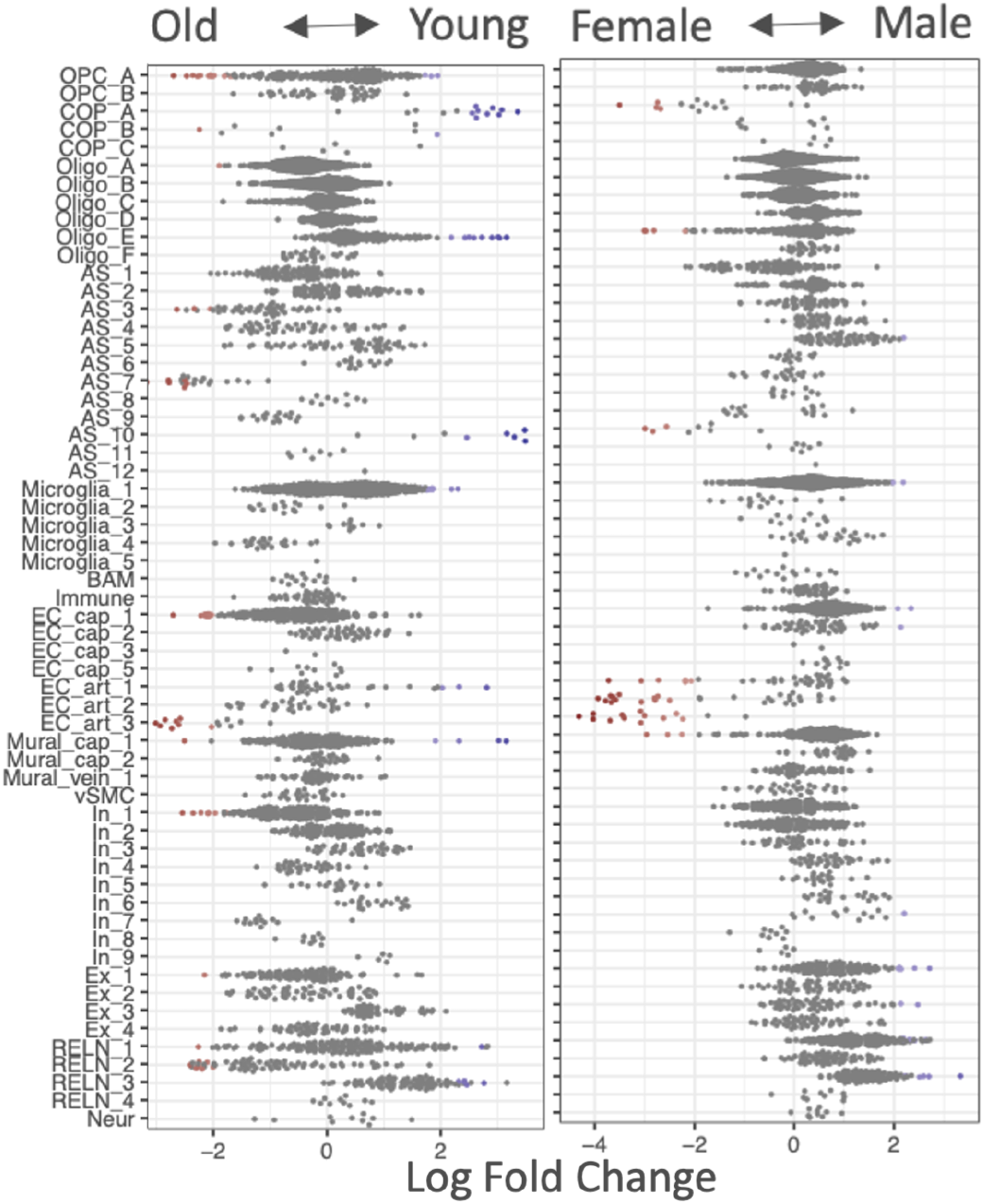
Compositional analysis between ages and sexes using Milo shows few differences.

## References

1. Bøstrand, S. M. K. & Williams, A. Oligodendroglial Heterogeneity in Neuropsychiatric Disease. Life 11, 125, DOI: 10.3390/life11020125 (2021).

2. Floriddia, E. M. et al. Distinct oligodendrocyte populations have spatial preference and different responses to spinal cord injury. Nat. Commun. 11, 1–15, DOI: 10.1038/s41467-020-19453-x (2020).

3. Marques, S. et al. Oligodendrocyte heterogeneity in the mouse juvenile and adult central nervous system. Science 352, 1326–1329, DOI: 10.1126/science.aaf6463 (2016).

4. Viganò, F., Möbius, W., Götz, M. & Dimou, L. Transplantation reveals regional differences in oligodendrocyte differentiation in the adult brain. Nat. Neurosci. 16, 1370–1372, DOI: 10.1038/nn.3503 (2013).

5. Bechler, M. E., Byrne, L., Bechler, M. E. & Byrne, L. CNS Myelin Sheath Lengths Are an Intrinsic Property of Oligodendrocytes. Curr. Biol. 25, 2411–2416, DOI: 10.1016/j.cub.2015.07.056 (2015).

6. Segel, M. et al. Niche stiffness underlies the ageing of central nervous system progenitor cells. Nature 573, 130–134, DOI: 10.1038/s41586-019-1484-9 (2019).

7. Crawford, A. H., Tripathi, R. B., Richardson, W. D. & Franklin, R. J. Developmental Origin of Oligodendrocyte Lineage Cells Determines Response to Demyelination and Susceptibility to Age-Associated Functional Decline. Cell Reports 15, 761–773, DOI: 10.1016/j.celrep.2016.03.069 (2016).

8. Leong, S. Y. et al. Heterogeneity of oligodendrocyte progenitor cells in adult human brain. Annals Clin. Transl. Neurol. 1, 272–283, DOI: 10.1002/acn3.55 (2014).

9. Spitzer, S. O. et al. Oligodendrocyte Progenitor Cells Become Regionally Diverse and Heterogeneous with Age. Neuron 101, 459–471, DOI: 10.1016/j.neuron.2018.12.020 (2019).

10. Neumann, B. et al. Metformin Restores CNS Remyelination Capacity by Rejuvenating Aged Stem Cells. Cell Stem Cell 25, 473–485, DOI: 10.1016/j.stem.2019.08.015 (2019).

11. Sim, F. J., Zhao, C., Penderis, J. & Franklin, R. J. M. The Age-Related Decrease in CNS Remyelination Efficiency Is Attributable to an Impairment of Both Oligodendrocyte Progenitor Recruitment and Differentiation. The J. Neurosci. 22, 2451–2459, DOI: 10.1523/JNEUROSCI.22-07-02451.2002 (2002).

12. Shobin, E. et al. Microglia activation and phagocytosis: relationship with aging and cognitive impairment in the rhesus monkey. GeroScience 39, 199–220, DOI: 10.1007/s11357-017-9965-y (2017).

13. Scalfari, A., Neuhaus, A., Daumer, M., Ebers, G. C. & Muraro, P. A. Age and disability accumulation in multiple sclerosis. Neurology 77, 1246–1252, DOI: 10.1212/WNL.0b013e318230a17d (2011).

14. Kalincik, T. et al. Sex as a determinant of relapse incidence and progressive course of multiple sclerosis. Brain 136, 3609–3617, DOI: 10.1093/brain/awt281 (2013).

15. Caceres, A., Jene, A., Esko, T., Perez-Jurado, L. A. & Gonzalez, J. R. Extreme downregulation of chromosome Y and Alzheimer’s disease in men. Neurobiol. Aging 90, 1–150, DOI: 10.1016/j.neurobiolaging.2020.02.003 (2020).

16. Chowen, J. A. & Garcia-Segura, L. M. Role of glial cells in the generation of sex differences in neurodegenerative diseases and brain aging. Mech. Ageing Dev. 196, 111473, DOI: 10.1016/j.mad.2021.111473 (2021).

17. Yasuda, K. et al. Sex-specific differences in transcriptomic profiles and cellular characteristics of oligodendrocyte precursor cells. Stem Cell Res. 46, 101866, DOI: 10.1016/j.scr.2020.101866 (2020).

18. Kodama, L. & Gan, L. Do Microglial Sex Differences Contribute to Sex Differences in Neurodegenerative Diseases? Trends Mol. Medicine 25, 741–749, DOI: 10.1016/j.molmed.2019.05.001 (2019).

19. Darmanis, S. et al. A survey of human brain transcriptome diversity at the single cell level. Proc. Natl. Acad. Sci. United States Am. 112, 7285–7290, DOI: 10.1073/pnas.1507125112 (2015).

20. Zhong, S. et al. A single-cell RNA-seq survey of the developmental landscape of the human prefrontal cortex. Nature 555, 524–528, DOI: 10.1038/nature25980 (2018).

21. Couturier, C. P. et al. Single-cell RNA-seq reveals that glioblastoma recapitulates a normal neurodevelopmental hierarchy. Nat. Commun. 11, DOI: 10.1038/s41467-020-17186-5 (2020).

22. Fu, Y. et al. Heterogeneity of glial progenitor cells during the neurogenesis-to-gliogenesis switch in the developing human cerebral cortex. Cell Reports 34, 108788, DOI: 10.1016/j.celrep.2021.108788 (2021).

23. Eze, U. C., Bhaduri, A., Haeussler, M., Nowakowski, T. J. & Kriegstein, A. R. Single-cell atlas of early human brain development highlights heterogeneity of human neuroepithelial cells and early radial glia. Nat. Neurosci. 24, 584–594, DOI: 10.1038/s41593-020-00794-1 (2021).

24. Huang, W. et al. Origins and Proliferative States of Human Oligodendrocyte Precursor Cells. Cell 182, 594–608, DOI: 10.1016/j.cell.2020.06.027 (2020).

25. Jäkel, S. et al. Altered human oligodendrocyte heterogeneity in multiple sclerosis. Nature 566, 543–547, DOI: 10.1038/s41586-019-0903-2 (2019).

26. Schirmer, L. et al. Neuronal vulnerability and multilineage diversity in multiple sclerosis. Nature 573, 75–82, DOI: 10.1038/s41586-019-1404-z (2019).

27. Mathys, H. et al. Single-cell transcriptomic analysis of Alzheimer’s disease. Nature 570, 332–337, DOI: 10.1038/s41586-019-1195-2 (2019).

28. Nagy, C. et al. Single-nucleus transcriptomics of the prefrontal cortex in major depressive disorder implicates oligodendrocyte precursor cells and excitatory neurons. Nat. Neurosci. 23, 771–781, DOI: 10.1038/s41593-020-0621-y (2020).

29. Yoshikawa, F. et al. Opalin, a Transmembrane Sialylglycoprotein Located in the Central Nervous System Myelin Paranodal Loop Membrane. J. Biol. Chem. 283, 20830–20840, DOI: 10.1074/jbc.M801314200 (2008).

30. Chang, K. J. et al. Glial ankyrins facilitate paranodal axoglial junction assembly. Nat. Neurosci. 17, 1673–1681, DOI: 10.1038/nn.3858 (2014).

31. Falcão, A. M. et al. Disease-specific oligodendrocyte lineage cells arise in multiple sclerosis. Nat. Medicine 24, 1837–1844, DOI: 10.1038/s41591-018-0236-y (2018).

32. Feldmann, A. et al. Transport of the major myelin proteolipid protein is directed by VAMP3 and VAMP7. J. Neurosci. 31, 5659–5672, DOI: 10.1523/JNEUROSCI.6638-10.2011 (2011).

33. Swire, M. et al. Oligodendrocyte hcn2 channels regulate myelin sheath length. J. Neurosci. 41, 7954–7964, DOI: 10.1523/JNEUROSCI.2463-20.2021 (2021).

34. van Bruggen, D. et al. Developmental landscape of human forebrain at a single-cell level unveils early waves of oligodendrogenesis. bioRxiv 2021.07.22.453317, DOI: 10.1101/2021.07.22.453317 (2021).

35. Saelens, W., Cannoodt, R., Todorov, H. & Saeys, Y. A comparison of single-cell trajectory inference methods. Nat. Biotechnol. 37, 547–554, DOI: 10.1038/s41587-019-0071-9 (2019).

36. Trapnell, C. et al. The dynamics and regulators of cell fate decisions are revealed by pseudotemporal ordering of single cells. Nat. Biotechnol. 32, 381–386, DOI: 10.1038/nbt.2859 (2014).

37. Street, K. et al. Slingshot: cell lineage and pseudotime inference for single-cell transcriptomics. BMC Genomics 19, 477, DOI: 10.1186/s12864-018-4772-0 (2018).

38. Bergen, V., Lange, M., Peidli, S., Wolf, F. A. & Theis, F. J. Generalizing RNA velocity to transient cell states through dynamical modeling. Nat. Biotechnol. 38, 1408–1414, DOI: 10.1038/s41587-020-0591-3 (2020).

39. Butler, A., Hoffman, P., Smibert, P., Papalexi, E. & Satija, R. Integrating single-cell transcriptomic data across different conditions, technologies, and species. Nat. Biotechnol. 36, 411–420, DOI: 10.1038/nbt.4096 (2018).

40. Sathyamurthy, A. et al. Massively Parallel Single Nucleus Transcriptional Profiling Defines Spinal Cord Neurons and Their Activity during Behavior. Cell Reports 22, 2216–2225, DOI: 10.1016/j.celrep.2018.02.003 (2018).

41. Gargareta, V.-I. et al. Conservation and divergence of myelin proteome and oligodendrocyte transcriptome profiles between humans and mice. bioRxiv 2022.01.17.476643 (2022).

42. Khakh, B. S. & Deneen, B. The Emerging Nature of Astrocyte Diversity. Annu. Rev. Neurosci. 42, 187–207, DOI: 10.1146/annurev-neuro-070918-050443 (2019).

43. Smith, A. M. et al. Diverse human astrocyte and microglial transcriptional responses to Alzheimer’s pathology. Acta Neuropathol. DOI: 10.1007/s00401-021-02372-6 (2021).

44. Al-Dalahmah, O. et al. Single-nucleus RNA-seq identifies Huntington disease astrocyte states. Acta Neuropathol. Commun. 8, 19, DOI: 10.1186/s40478-020-0880-6 (2020).

45. Zhou, Y. et al. Human and mouse single-nucleus transcriptomics reveal TREM2-dependent and TREM2-independent cellular responses in Alzheimer’s disease. Nat. Medicine 26, 131–142, DOI: 10.1038/s41591-019-0695-9 (2020).

46. Sharma, A. et al. Single-cell atlas of progressive supranuclear palsy reveals a distinct hybrid glial cell population. bioRxiv 2021.04.11.439393 (2021).

47. Sterpka, A. & Chen, X. Neuronal and astrocytic primary cilia in the mature brain. Pharmacol. Res. 137, 114–121, DOI: 10.1016/j.phrs.2018.10.002 (2018).

48. Zamanian, J. L. et al. Genomic analysis of reactive astrogliosis. J. Neurosci. 32, 6391–6410, DOI: 10.1523/JNEUROSCI.6221-11.2012 (2012).

49. Liddelow, S. A. et al. Neurotoxic reactive astrocytes are induced by activated microglia. Nature 541, 481–487, DOI: 10.1038/nature21029 (2017).

50. Moulson, A. J., Squair, J. W., Franklin, R. J., Tetzlaff, W. & Assinck, P. Diversity of Reactive Astrogliosis in CNS Pathology: Heterogeneity or Plasticity? Front. Cell. Neurosci. 15, DOI: 10.3389/fncel.2021.703810 (2021).

51. Escartin, C. et al. Reactive astrocyte nomenclature, definitions, and future directions. Nat. Neurosci. 24, 312–325, DOI: 10.1038/s41593-020-00783-4 (2021).

52. Clarke, L. E. et al. Normal aging induces A1-like astrocyte reactivity. Proc. Natl. Acad. Sci. United States Am. 115, E1896–E1905, DOI: 10.1073/pnas.1800165115 (2018).

53. Kreutzberg, G. W. Microglia: A sensor for pathological events in the CNS. Trends Neurosci. 19, 312–318, DOI: 10.1016/0166-2236(96)10049-7 (1996).

54. Hanisch, U. K. & Kettenmann, H. Microglia: Active sensor and versatile effector cells in the normal and pathologic brain. Nat. Neurosci. 10, 1387–1394, DOI: 10.1038/nn1997 (2007).

55. Kierdorf, K. & Prinz, M. Microglia in steady state. J. Clin. Investig. 127, 3201–3209, DOI: 10.1172/JCI90602 (2017).

56. Reemst, K., Noctor, S. C., Lucassen, P. J. & Hol, E. M. The Indispensable Roles of Microglia and Astrocytes during Brain Development. Front. Hum. Neurosci. 10, 1–28, DOI: 10.3389/fnhum.2016.00566 (2016).

57. Dann, E., Henderson, N. C., Teichmann, S. A., Morgan, M. D. & Marioni, J. C. Differential abundance testing on single-cell data using k-nearest neighbor graphs. Nat. Biotechnol. DOI: 10.1038/s41587-021-01033-z (2021).

58. Pringle, N. P. & Richardson, W. D. A singularity of PDGF alpha-receptor expression in the dorsoventral axis of the neural tube may define the origin of the oligodendrocyte lineage. Development 117, 525–533, DOI: 10.1242/dev.117.2.525 (1993).

59. Lake, B. B. et al. Integrative single-cell analysis of transcriptional and epigenetic states in the human adult brain. Nat. Biotechnol. 36, 70–80, DOI: 10.1038/nbt.4038 (2018).

60. Khandker, L. et al. Single-cell Sequencing Reveals Brain/Spinal Cord Oligodendrocyte Precursor Heterogeneity and Requirement for mTOR in Cholesterol Biosynthesis and Myelin Maintenance. bioRxiv 2020.10.22.349209, DOI: 10.1101/2020.10.22.349209 (2020).

61. Ghelman, J. et al. SKAP2 as a new regulator of oligodendroglial migration and myelin sheath formation. Glia 69, 2699–2716, DOI: 10.1002/glia.24066 (2021).

62. Yoon, H., Walters, G., Paulsen, A. R. & Scarisbrick, I. A. Astrocyte heterogeneity across the brain and spinal cord occurs developmentally, in adulthood and in response to demyelination. PLOS ONE 12, e0180697, DOI: 10.1371/journal.pone.0180697 (2017).

63. Bugiani, M., Plug, B. C., Man, J. H., Breur, M. & van der Knaap, M. S. Heterogeneity of white matter astrocytes in the human brain. Acta Neuropathol. DOI: 10.1007/s00401-021-02391-3 (2021).

64. Gerrits, E. et al. Distinct amyloid-*β* and tau-associated microglia profiles in Alzheimer’s disease. Acta Neuropathol. 141, 681–696, DOI: 10.1007/s00401-021-02263-w (2021).

65. Moruzzo, D. et al. The Transcription Factors EBF1 and EBF2 Are Positive Regulators of Myelination in Schwann Cells. Mol. Neurobiol. 54, 8117–8127, DOI: 10.1007/s12035-016-0296-2 (2017).

66. Prasad, B. et al. unc-3, a gene required for axonal guidance in Caenorhabditis elegans, encodes a member of the O/E family of transcription factors. Development 125, 1561–1568, DOI: 10.1242/dev.125.8.1561 (1998).

67. Martínez, A. et al. Early B-cell Factor gene association with multiple sclerosis in the Spanish population. BMC Neurol. 5, 19, DOI: 10.1186/1471-2377-5-19 (2005).

68. Favuzzi, E. et al. Activity-Dependent Gating of Parvalbumin Interneuron Function by the Perineuronal Net Protein Brevican. Neuron 95, 639–655, DOI: 10.1016/j.neuron.2017.06.028 (2017).

69. Thrupp, N. et al. Single-Nucleus RNA-Seq Is Not Suitable for Detection of Microglial Activation Genes in Humans. Cell Reports 32, 108189, DOI: 10.1016/j.celrep.2020.108189 (2020).

70. Alsema, A. M. et al. Profiling Microglia From Alzheimer’s Disease Donors and Non-demented Elderly in Acute Human Postmortem Cortical Tissue. Front. Mol. Neurosci. 13, 1–14, DOI: 10.3389/fnmol.2020.00134 (2020).

71. Fernández, M. V. et al. Evaluation of gene-based family-based methods to detect novel genes associated with familial late onset Alzheimer disease. Front. Neurosci. 12, DOI: 10.3389/fnins.2018.00209 (2018).

72. Wang, X. et al. DUSP1 Promotes Microglial Polarization toward M2 Phenotype in the Medial Prefrontal Cortex of Neuropathic Pain Rats via Inhibition of MAPK Pathway. ACS Chem. Neurosci. 12, 966–978, DOI: 10.1021/acschemneuro.0c00567 (2021).

73. Bramow, S. et al. Demyelination versus remyelination in progressive multiple sclerosis. Brain 133, 2983–2998, DOI: 10.1093/brain/awq250 (2010).

74. Neumann, B., Segel, M., Chalut, K. J. & Franklin, R. J. Remyelination and ageing: Reversing the ravages of time. Multiple Scler. J. 25, 1835–1841, DOI: 10.1177/1352458519884006 (2019).

75. Yeung, M. S. et al. Dynamics of oligodendrocyte generation in multiple sclerosis. Nature 566, 538–542, DOI: 10.1038/s41586-018-0842-3 (2019).

76. Franklin, R. J. M. & Ffrench-Constant, C. Remyelination in the CNS: from biology to therapy. Nat. Rev. Neurosci. 9, 839–855, DOI: 10.1038/nrn2480 (2008).

77. Neely, S. et al. New oligodendrocytes exhibit more abundant and accurate myelin regeneration than those that survive demyelination. 1–14, DOI: 10.1101/2020.05.22.110551 (2020).

78. Bacmeister, C. M. et al. Motor learning promotes remyelination via new and surviving oligodendrocytes. Nat. Neurosci. 23, 819–831, DOI: 10.1038/s41593-020-0637-3 (2020).

79. McDavid, A., Finak, G. & Yajima, M. MAST: Model-based Analysis of Single Cell Transcriptomics. (2020).

80. Rue-Albrecht, K., Lun, A., Soneson, C. & Stadler, M. velociraptor: Toolkit for Single-Cell Velocity (2021).

81. La Manno, G. et al. RNA velocity of single cells. Nature 560, 494–498, DOI: 10.1038/s41586-018-0414-6 (2018).

82. Yu, G., Wang, L. G., Han, Y. & He, Q. Y. ClusterProfiler: An R package for comparing biological themes among gene clusters. OMICS A J. Integr. Biol. 16, 284–287, DOI: 10.1089/omi.2011.0118 (2012).

83. Bankhead, P. et al. QuPath: Open source software for digital pathology image analysis. Sci. Reports 7, 1–7, DOI: 10.1038/s41598-017-17204-5 (2017).

84. Sanmarco, L. M. et al. Gut-licensed IFN*γ*+ NK cells drive LAMP1+TRAIL+ anti-inflammatory astrocytes. Nature 590, 473–479, DOI: 10.1038/s41586-020-03116-4 (2021).

